# Basolateral amygdala circuits supporting negative emotional bias in a mouse model for depression

**DOI:** 10.1101/2023.01.23.525169

**Authors:** Mathilde Bigot, Claire-Hélène De Badts, Axel Benchetrit, Éléonore Vicq, Carine Moigneu, Manon Meyrel, Sébastien Wagner, Josselin Houenou, Pierre-Marie Lledo, Chantal Henry, Mariana Alonso

## Abstract

Negative emotional bias is an essential hallmark of depression reflected by negative shift in hedonic valence assignment to emotional stimuli. Pleasant cues become less attractive and unpleasant ones more aversive. Given the crucial role of amygdala in valence coding, we hypothesize that specific basolateral amygdala (BLA) circuits alterations might support negative emotional bias associated with depressive states.

Using a translational assay, we evaluate odor valence assignment in an animal model for depression chronically administered by corticosterone (CORT). We show spontaneous negative bias in depressive-like mice that attribute more negative valences for both attractive and aversive odors, mimicking thus the bias observed in depressed bipolar patients.

Combining CTB and rabies-based tracing with *ex vivo* measurements of neuronal activity and chemogenetics experiments, we find that the CORT treatment reduces BLA-to-nucleus accumbens (NAc) neuronal activity and increases BLA-to-central amygdala activity, circuits respectively known to be involved in positive and negative valence encoding. Alterations in presynaptic connectivity of BLA-projecting neurons accompany these activity shifts. Interestingly, inputs from the paraventricular thalamus nucleus (PVT) towards BLA-to-NAc neurons are reduced in CORT-treated mice. Finally, chemogenetically activating the BLA-to-NAc circuit attenuates the negative bias in CORT-treated mice as well as the depressive-like phenotype, similarly than Fluoxetine antidepressant treatment. Altogether, we demonstrate that depressive states are associated with negative emotional bias both in human and mice. This bias is supported by activity shifts of specific BLA circuits along with durable presynaptic connectivity changes, but it could be alleviated by antidepressant drug or activity manipulation of altered BLA circuit.

## Introduction

Depression is the single largest contributor to disability worldwide, affecting 300 million people annually. Depressive episodes occur in patients suffering from major depressive disorder (MDD) and bipolar disorders (BD). Negative emotional bias is an essential characteristic of depressive episodes leading patients to attribute more negative valence to salient environmental cues (1–4). Restoration of these emotional biases is predictive of the response to antidepressant treatment and essential for recovery from the episode (5,6). Importantly, emotional biases can be explored in animals using approach or avoidance behaviors which are quantifiable motor readouts indicating the valence assigned to stimuli (7–9).

In humans, numerous studies have shown the amygdala involvement in the etiopathology of depression with changes in volume, connectivity, and activity responses to emotional tasks (10). Recently, robust findings in animals have highlighted how basolateral amygdala (BLA) circuits assign positive and negative valences to emotional stimuli. During associative learning, BLA neurons preferentially involved either in positive or negative valence coding can be distinguished according to their projection targets as well as spatial and genetic characteristics (11). The BLA neurons projecting to the nucleus accumbens (NAc) mainly respond to positive stimuli and trigger approach behaviors, while BLA neurons targeting the centromedial nucleus of the amygdala (CeA) mainly respond to negative stimuli, generating defensive responses, despite genetic and functional heterogeneity reported in some studies (12–16).

How depressive states alter valence assignment is still unknown. Here, we developed a translational test to reveal the presence of spontaneous negative emotional bias in a mouse model for depression, as we observed in depressed BD patients. We show that depressive-like states are associated with specific BLA circuits activity disturbances, and that manipulating such circuits attenuates the negative bias.

## Methods and Materials

### Human subjects and Sniffin’ sticks test

The Human Research Ethics Committee, CPP-Ile de France IV (2015/44), approved the study and all participants gave their informed consent. Forty-eight patients were included in our study, aged from 18–65 years, with a current BD diagnosis according to the DSM-IV-R criteria (17) and 11 healthy subjects. Depression severity was measured by MADRS scale.

Odor valence assignment was evaluated during the identification task of the Sniffin’ sticks test (Burghardt®, Germany), as previously published (18).

### Animals and chronic corticosterone administration model for depression

All animal care and experimental procedures followed national and European (2010/63/EU) guidelines and were approved by the French Ministry of Research (APAFiS: #16380-2018080217358599_v1). C57BL/6N male mice (7–8 weeks old, Taconic Farms (Denmark), n = 181) were socially housed (4-6 per cage) and maintained under standard conditions (23 ± 1 °C; humidity 40%) on a 14/10 h light/dark cycle with food and water *ad libitum*.

Corticosterone (CORT, 35 μg/ml, equivalent to about 5 mg/kg/day) was prepared in vehicle made of 0.45% (2-hydroxypropyl)-beta-cyclodextrin (β-CD; both from Sigma-Aldrich, France), and administered in drinking water for 28 days, as previously described (19). After four weeks of CORT treatment, the fluoxetine hydrochloride antidepressant (equivalent to about 18 mg/kg/day; Sigma-Aldrich) was administered for three weeks along with CORT administration (19).

### Behavioral assessment

Several behavioral tests assessed anxiety, depression-like phenotypes, and olfactory preferences. The olfactory preference test was performed in a quiet and dimly lit room (adapted from (20)). Clean housing cages with regular bedding material were used as testing arenas. A petri dish with hooled cover was placed at one side of the arena. The behavior was recorded by video camera for 15 min and the Noldus Ethovision 3.0 (Netherlands) system was used to track position and locomotion of mice. During the first 4 days, only a Whatman paper filter (GE Healthcare Life Sciences, USA) was placed into the petri dish, to assess baseline exploration. Then, 2 days were dedicated to each odor with the odorant placed on paper filter in the following order: peanut oil (pure, 400 ul), female urine (pure, 100 ul), trimethylamine (Sigma-Aldrich, 6.75% in water, 400 ul) and trimethylthiazoline (Sigma-Aldrich, 5% in mineral oil, 400 ul).

### CTB stereotaxic injections

To label neurons of the BLA projecting to NAc or CeA, the retrograde tracer Cholera Toxin Subunit B conjugated with Alexa Fluor 555 or 674 (CTB 555 or CTB 647, 1 mg/ml, Invitrogen, USA) was bilaterally injected with a micropipette connected to Nanoject III microinjector (Drummond Scientific) in the NAc (AP: +1.4 mm; ML: ±0.87 mm; DV: −4.35 mm, 300 nl) or in the CeA (AP: −0.8 mm, ML: ±2.35mm, DV: −4.35 mm, 100 nl, targeting the centromedial part).

### Viral stereotaxic injections

For labelling presynaptic inputs to BLA-to-NAc and BLA-to-CeA neurons, AAV-FLEX-G-TVA-GFP was injected bilaterally in the BLA (AP: −1.75 mm, ML: ±3.15 mm, DV: −4.25 mm, 100 nl) and AAVretro-CRE either in the NAc (100 nl) or in the CeA (50 nl) using previously mentioned coordinates. After four weeks of CORT treatment, mice were injected with EnvA-RVΔG-mCherry (150 nl) in the BLA and sacrificed one week later.

AAV5-hSyn-DIO-hM3Dq-mCherry, AAV5-hSyn-DIO-mCherry and AAVretro-PGK-Cre recombinant adeno-associated virus were used to manipulate BLA neuronal activity. Viral vectors were bilaterally injected into the BLA (100-150 nl) and the NAc (100-150 nl) or the CeA (50 nl) using previously mentioned coordinates.

All AAV were provided by Addgene (USA) and EnvA-RVΔG-mCherry was generated by Karl-Klaus Conzelmann lab (University of Munich, Germany).

### Immunohistochemistry and image analysis

Mice were anesthetized with ketamine/xylazine mix (intraperitoneal (i.p)., 150 mg/kg and 50 mg/kg respectively) and perfused transcardially with 0.9% NaCl, followed by 4% paraformaldehyde in phosphate buffer, pH 7.3. Forty-micrometer-thick coronal brain sections were obtained using a sliding microtome (Leica SM 2010 R). Immunostaining was performed on free-floating sections using the following antibodies: guinea pig anti-cFos (1:2000, Synaptic Systems, Germany), rabbit anti-RFP (1:4000, Rockland, USA), chicken anti-GFP (1:1000, Abcam), and secondary antibodies (Alexa-conjugated secondary antibodies at 1:1000, Jackson ImmunoResearch Laboratories, United Kindom and Molecular Probes, USA).

### Statistical analysis

Statistical analyses were performed with GraphPad Prism v9 software (USA). Two-sided parametric or non-parametric tests were used according to the normality of datasets and variances homoscedasticity. Following post-hoc analyses were applied with the False Rate Discovery (FDR) correction method.

All datasets were described using the mean; error bars in the figures represent standard error mean (SEM), except for **Figure S5E-G** where box-and-whiskers plots were used (boxes represent the 25^th^, median and 75^th^ percentiles while whiskers represent the 1^st^ and 99^th^ percentiles). Differences were considered significant for p < 0.05.

The datasets and codes that support the findings of this study are available from the corresponding author upon reasonable request.

Detailed methods and materials can be found in the Supplemental information.

## Results

### Negative olfactory valence bias in depressed BD patients

We evaluated olfactory performance using the Sniffin’ sticks test in a cohort of BD patients compared to control subjects. Patients were classified based on the DSM-IV-R (17) as euthymic, and depressed or mixed (referred to collectively as “depressed BD patients”). The group of depressed BD patients had significantly higher MADRS score, measuring depression severity, than both control subjects and euthymic BD patients, while euthymic BD patients did not differ from control subjects (**Table 1**). Further demographic and clinical characteristics are presented in **Table 1**.

**Table 1.** Demographic and clinical characteristics of BD patients and control subjects. BD NOS: bipolar disorder not otherwise specified; MADRS: Montgomery Asberg Depression Rating Scale; YMRS: Young Mania Rating Scale.

| Variable | Control subjects (n = 11) | Euthymic BD patients (n = 25) | Depressed mixed BD patients (n = 23) | F, H or $\chi^2$ | p |
| --- | --- | --- | --- | --- | --- |
| Female, n (%) | 7 (63.63) | 13 (52) | 12 (52.17) | 0.48 | 0.786 |
| Age, years mean (SD) | 33.55 (10.8) | 38.2 (12.55) | 43.09 (11.33) | 2.61 | 0.083 |
| Education, years mean (SD) | 16 (3.03) | 14.65 (2.48) | 15.86 (3.51) | 1.21 | 0.306 |
| Unemployed, n (%) | 1 (9.09) | 11 (44) | 7 (31.82) | 4.24 | 0.12 |
| Current smoking, n (%) | 2 (18.18) | 12 (48) | 7 (30.43) | 3.4 | 0.183 |
| Type BD, n (%) |  |  |  |  |  |
| BD I |  | 16 (64) | 13 (56.52) |  |  |
| BD II |  | 9 (36) | 9 (39.13) |  |  |
| BD NOS |  | 0 (0) | 1 (4.35) |  |  |
| MADRS, mean (SD) | 1.36 (2.38) | 4.92 (5.82) | 22.61 (6.97) | 41.15 | < 0.001 |
| Control subjects vs Euthymic BD patients |  |  |  |  | 0.339 |
| Control subjects vs Depressed BD patients |  |  |  |  | < 0.001 |
| Euthymic BD patients vs Depressed BD patients |  |  |  |  | < 0.001 |
| YMRS | 0.36 (0.81) | 1.12 (2.57) | 3.83 (5.46) | 12.26 | 0.002 |
| Control subjects vs Euthymic BD patients |  |  |  |  | 0.459 |
| Control subjects vs Depressed BD patients |  |  |  |  | 0.003 |
| Euthymic BD patients vs Depressed BD patients |  |  |  |  | 0.004 |

The Sniffin’ sticks test (18,21,22) evaluates odor detection, discrimination, identification and valence (**Figure 1A**). Control subjects and all BD patients had similar detection, discrimination and identification scores (**Figure 1B-D**), suggesting no olfactory dysfunction in BD patients. However, depressed BD patients rated less odors as pleasant than euthymic BD patients, and more odors as unpleasant than both control subjects and euthymic BD patients (**Figure 1E**). This negative olfactory valence bias was associated with depression severity, as the MADRS score correlated negatively with the number of classified pleasant odors, and positively with the number of classified unpleasant odors in BD patients (**Figure 1F**). No correlation was found between the number of classified neutral odors and depression severity (**Figure 1F**).

**Figure 1.**
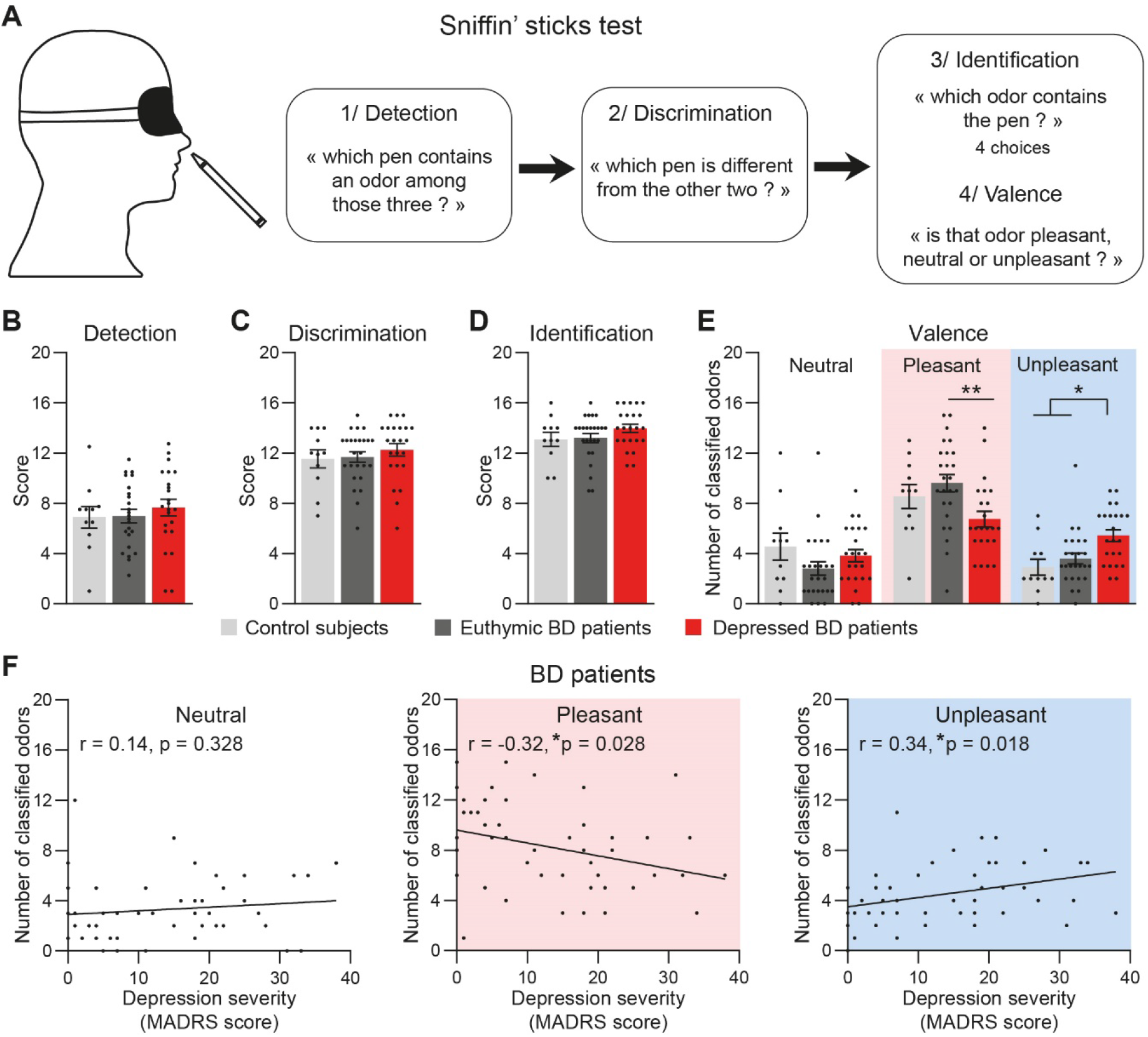
Negative olfactory valence bias in depressed BD patients. (**A**), Scheme of the Sniffin’ sticks test evaluating olfactory detection, discrimination and identification. Each of the presented odors for identification were then classified among neutral, pleasant or unpleasant valence. (**B-D**), BD patients (euthymic, n = 25 (except in (**B**): n = 24); depressed, n = 23) and control subjects (n = 11) did not differ on olfactory performances for detection (**B**, F(2,55) = 0.42, p = 0.659), discrimination (**C**, F(2,56) = 0.54, p = 0.586), and identification (**D**, H = 2.26, p = 0.323). (**E**), Depressed BD patients classified less odors as pleasant ones than euthymic BD patients (**p < 0.01), and more as unpleasant ones compared to euthymic BD patients and control subjects (*p < 0.05) (Group: F(2,56) = 0.00, p > 0.999, Valence: F(2,112) = 30.1, p < 0.001, Interaction: F(4,112) = 4.15, p = 0.004). (**F**) BD patients with higher depression severity (*i.e*. higher MADRS score) classified less odors as pleasant (middle), and more odors as unpleasant (right). The number of classified neutral odors did not change with depression severity (left). Bars are mean ± sem.

### Negative olfactory valence bias in a CORT-induced mouse model for depression

Then, negative olfactory valence bias was evaluated in mouse model for depression. We chose the well-described mouse model for depression induced by four weeks of CORT administration in drinking water (19) (**Figure 2A** and **Figure S1A-B**). We confirmed the anxiety-like phenotype in the open field (OF) and light and dark box (LDb) (**Figure 2B** and **Figure S1C-G**), and the depressive-like behavior in the splash test (ST) (**Figure 2B** and **Figure S1H-J**; z-scores calculated with the different parameters recorded for each behavioral test as described in Supplementary information). As previously reported, CORT-treated mice did not express despair-like features in the tail suspension test (TST) (23) (**Figure 2B** and **Figure S1K-M**). Overall, global emotionality score was significantly increased in CORT mice confirming an anxiety and depressive-like phenotype (**Figure 2C**).

**Figure 2.**
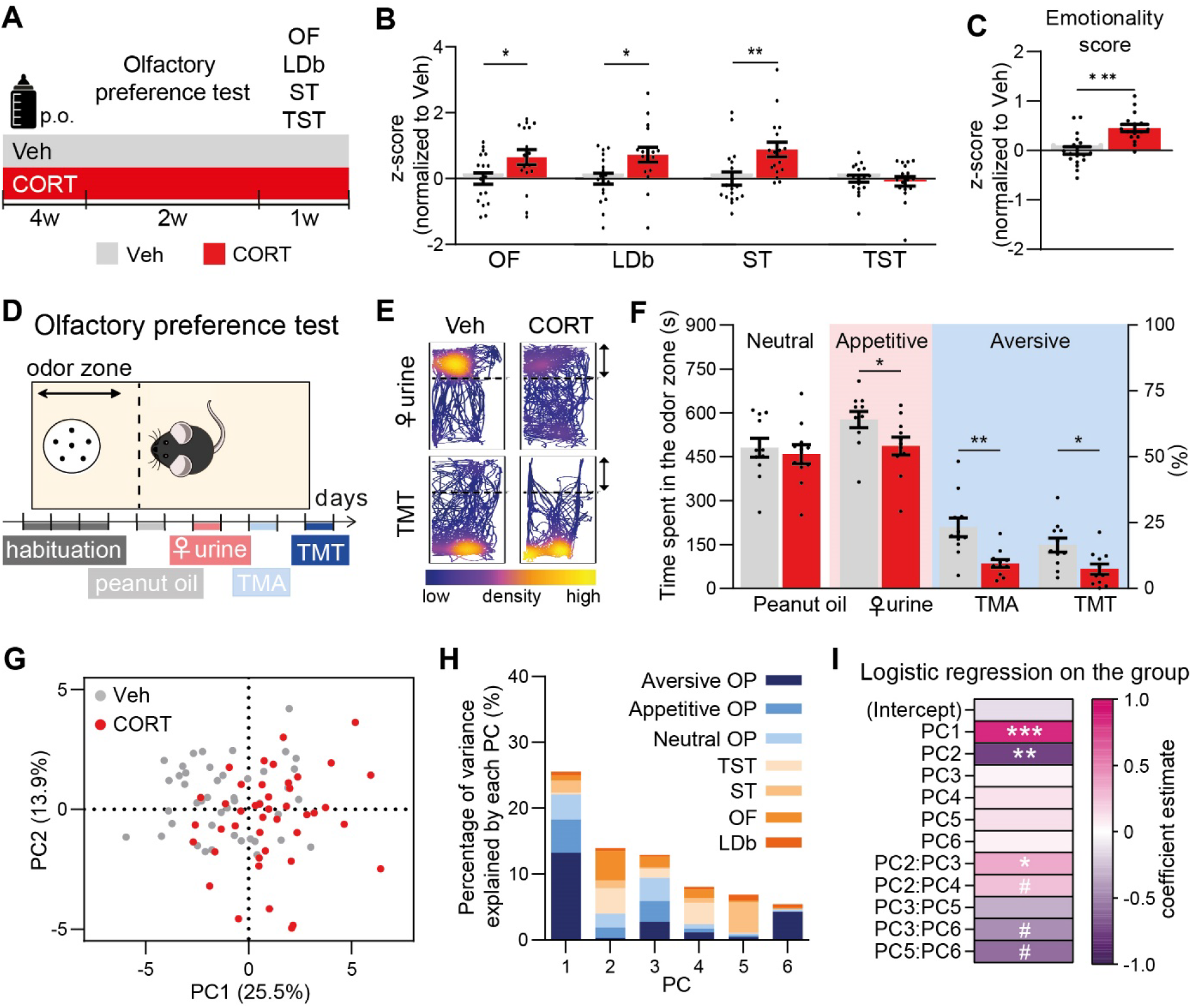
Negative olfactory valence bias in mouse depression model. (**A**) Mice received vehicle (grey, n = 18) or chronic CORT (red, n = 17) to model depression. (**B**) CORT mice presented anxiety-like phenotypes in the open field (OF, t(33) = 2.26, p = 0.030) and light and dark box (LDb, t(33) = 2.62, p = 0.013), and depressive-like behaviors in the splash test (ST, U = 59, p = 0.001) but not in the tail suspension test (TST, U = 149, p = 0.909). (**C**) Global emotionality score is increased in CORT mice (t(32) = 4.23). (**D**) Scheme of the olfactory preference test using neutral (grey), appetitive (pink) and aversive (blue) odors. (**E**) Representative mouse tracks colored by the density of position points. (**F**) CORT mice explored less ♀ urine, TMA and TMT than Veh controls (Group: F(1,20) = 10.63, p = 0.004; Odor: F(3.60) = 163.50, p < 0.001; Interaction: F(3.60) = 1.54, p = 0.214). (**G**) Principal Component Analysis on the 23 behavioral parameters evaluated in 87 mice from both CORT (red) and Veh (grey) mice. (**H**) The olfactory preference test (OP) standed for 86% of contributions to PC1 (blue bar parts). The OF (32%) and TST (28%) contributed mostly to PC2. (**I**) PC1, PC2 and PC2:PC3 significantly predicted the Veh/CORT status. Bars are mean ± sem. ^#^0.05 ≤ p < 0.1, *p < 0.05, **p < 0.01, ***p < 0.001. Bars are mean ± sem.

To assess valence assignment in mice, we set up an olfactory preference test (**Figure 2D**). Each odor was presented on two consecutive days, and its attributed valence was defined by increased, decreased or similar odor zone exploration time compared to habituation in Vehicle (Veh)-treated controls (One-way repeated measures (RM) ANOVA: F(4,40) = 44.86, p < 0.001, followed by FDR post-hoc comparisons). Peanut oil was neutral in our experimental conditions (t(40) = 1.36, q = 0.183), whereas ♀ urine was appetitive (t(40) = 3.85, q < 0.001) and synthetic compounds trimethylamine (TMA) and 2,4,5-trimethylthiazole (TMT) were aversive (TMA: t(40) = 5.68, q < 0.001, TMT: t(40) = 7.27, q < 0.001).

CORT-treated mice investigated both appetitive ♀ urine and aversive TMA and TMT in a lesser extent than Veh-treated controls, as revealed by decreased exploration time of the odor zone (**Figure 2E, F**) and farther mean location to the odor zone (**Figure S2E**). This negative olfactory valence bias was specific for non-neutral odors, without differences in the peanut oil exploration (**Figure 2F**), or during habituation (**Figure S2A-C**).

We wondered if this olfactory valence bias distinguishing CORT-treated mice from Veh-treated controls revealed variability already measured by other behavioral tests, or could be considered as new behavioral feature. Therefore, we applied principal component analysis (PCA) on a set of data containing 87 mice (44 Veh and 43 CORT) evaluated with all the tests presented in **Figure 2A**, *i.e*. through 23 behavioral parameters (**Figure 2G**). We selected the six first PCs, representing 72.6% of explained variance. Among each PC, we summed for each test the contribution of the different recorded behavioral parameters (**Figure 2H**), so that the neutral, appetitive and aversive parts of the olfactory preference test (OP) contributed respectively to 15%, 20% and 51% to the PC1, 27%, 24% and 21% to PC3, and 7%, <1% and 78% to PC6. Contributions to PC2 were mainly driven by the OF (32%) and the TST (28%). The TST was also the major contributor to PC4 (40%). The ST contributed to 7% of PC1 and 8% of PC2, but 68% of PC5, and the LDb contributed to 13% of PC5. The relative segregation of olfactory preference test measurements from other tests in the PC1, PC3 and PC6 suggested that these olfactory parameters did not correlate with the LDb, OF, ST and TST, but rather captured variability on another dimension. We then performed a logistic regression on the group (Veh/CORT status) using the PCs and their second order interactions as predictors. AIC-based stepwise regression allowed us to select the minimal number of predictors while keeping the most information (Supplemental information, **Figure 2I**). Predictors significantly associated with the group were PC1, PC2 and the interaction PC2:PC3, whereas PC2:PC4, PC3:PC6 and PC5:PC6 showed a statistical trend to association with the group. In summary, these results suggest that olfactory valence assignment is an independent and suitable behavioral measurement to predict differences between control and depressive-like states in mice.

### Fluoxetine improves negative emotional bias in responsive CORT treated-mice

Fluoxetine (FLX), a selective serotonin reuptake inhibitor, is a very commonly prescribed antidepressant drug. After four weeks of CORT administration, mice were treated with FLX in drinking water for three additional weeks. We compared treated animals with mice administered with CORT alone in the different behavioral tests used previously (**Figure 3A** and **Figure S3**). In agreement with previous data (24), FLX-treated group showed a significally reduced emotionality score compared to CORT group indicating reduced anxiety and depression levels, even though this group is extremely heterogeneous (**Figure S3D**). Clinical response to antidepressants in depression is defined as 50% decrease in rating scale score (25). Thus, we analyzed our behavioral data according to this 50% cut-off, as previously described (24). We defined FLX responsive mice with the criterion of global emotionality score under −0.5 (CORT+FLX-R) where mice with emotionality score above −0.5 were defined as non-responsive (CORT+FLX-NR; **Figure 3B**). Interestingly, CORT+FLX-R mice showed an increased exploration time for neutral and appetitive odors compared to CORT and CORT-FLX-NR mice (**Figure 3C, D**), and their mean position was closer to the odor zone for these stimuli (**Figure S3R**). Surprinsingly, TMA was more repulsive for CORT-FLX-NR mice, while there was no difference in TMT exploration compared to CORT mice (**Figure 3C, D** and **Figure S3R**).

**Figure 3.**
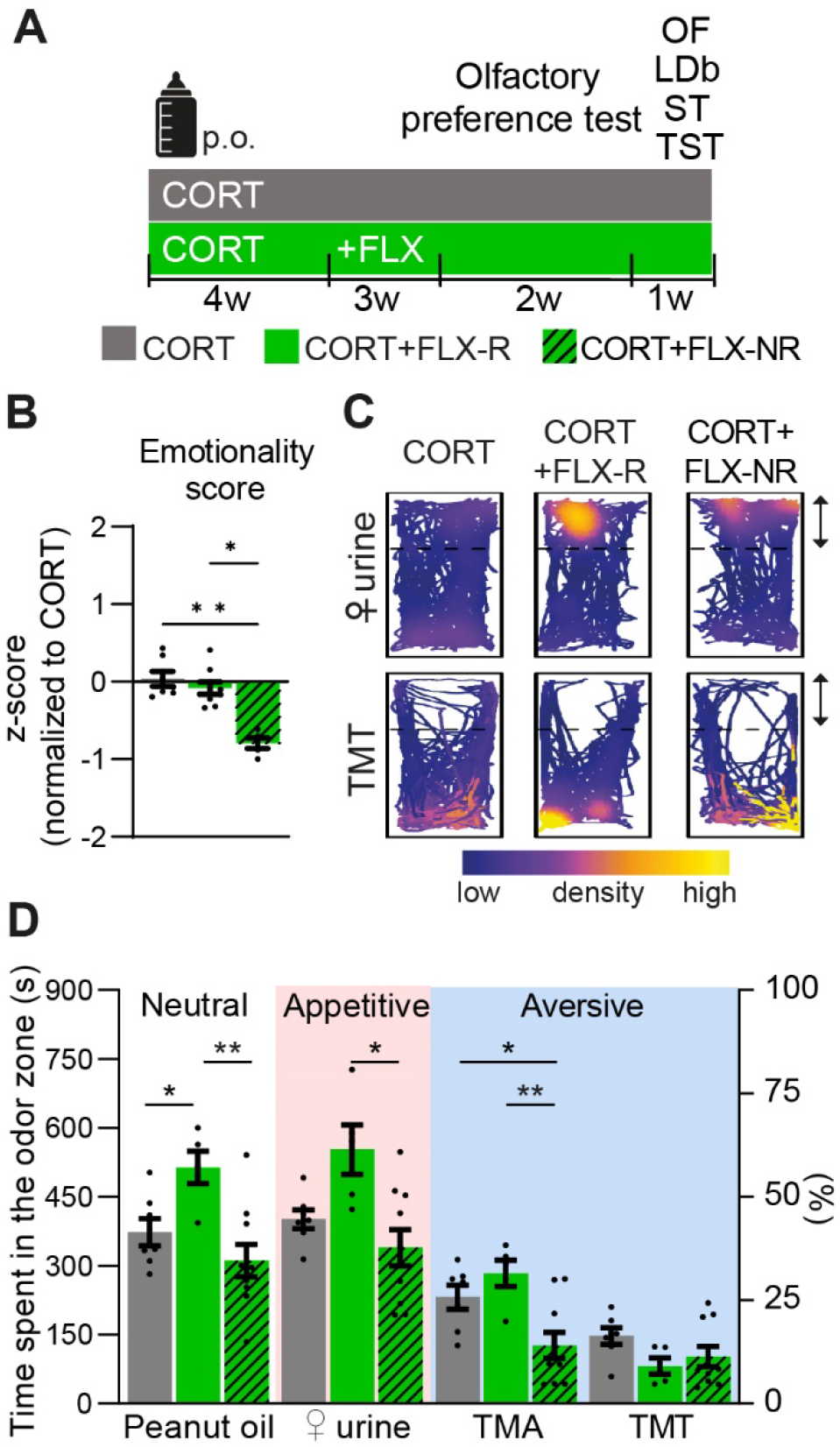
Fluoxetine partially reversed the CORT phenotype in responsive mice. (**A**) Mice received chronic CORT to model depression (grey, n = 7), or CORT for 4 weeks then fluoxetine in addition to CORT for the following weeks (green, n = 14). (**B**) The reduction of the global emotionality score shows an improvement of the depressed- and anxious-like phenotype in responsive mice (U = 11.78, p = 0.0005). Fluoxetine-responsive mice are defined by an emotionality score lower than −0.5. (**C**) Representative mouse tracks colored by the density of position points. (**D**) CORT+FLX-R mice explored more peanut oil and ♀ urine, but less TMA, than CORT and CORT+FLX-NR groups (Group: F(2,19) = 9.629, p = 0.0013; Odor: F(2.192) = 86.61, p < 0.001; Interaction: F(6.57) = 4.11, p = 0.0017). *p < 0.05, **p < 0.01. Bars are mean ± sem.

Importantly, no effect was observed during habituation or for distance moved (**Figure S3P-Q)**. These data demonstrate that restoration of depressive-like phenotype in responsive mice comes with partial improvement of negative emotional bias, specifically increasing attractiveness of neutral and pleasant odors.

### BLA circuits activity alterations in CORT-induced mouse model for depression

According to the major role of BLA in valence processing, we hypothesized that activity of BLA-to-NAc projecting neurons would be reduced in depressive states while BLA-to-CeA neurons would be more active. Thus, we injected retrograde CTB647 and CTB555 dyes in the NAc and CeA respectively (**Figure 4A** and **Figure S4**) to label BLA projecting neurons to these structures. Injections were performed in both CORT or Veh-treated mice, one week before presenting odors to trigger cFos expression, used as immunohistological proxy for neuronal activation (**Figure 4B**). We used Icy software to automatically detect cFos+, colocalizing cFos+/CTB647+, cFos+/CTB555+ and cFos+/CTB647+/CTB555+ cells (**Figure S5D-G**, Supplemental information). Although odorless mineral oil, appetitive ♀ urine and aversive TMT recruited different proportion of the BLA-to-NAc, BLA-to-CeA and BLA-to-NAc-and-CeA neurons both in Veh-treated control and in CORT-treated mice, the total density of cFos+ and colocalizing cFos+/CTB+ cells were mostly similar across odors and groups (**Figure S5H-J** and **Table S1**) and the total cFos+ cell number did not differ between Veh and CORT-treated groups in the BLA (**Figure 4D**). Interestingly, when compared regardless of the odor used for cFos stimulation, BLA-to-NAc neurons were less active in CORT-treated mice compared to Veh-treated controls, while BLA-to-CeA neurons more active (**Figure 4E, F**). BLA cells projecting to both NAc and CeA tended to be less recruited in CORT-treated mice than in Veh-treated controls (**Figure 4G**). Among all BLA cFos+ cells, the numbers of projecting neurons were differently distributed between groups, with less BLA-to-NAc cFos+ cells, and more BLA-to-CeA and unidentified cFos+ cells in the CORT-treated mice relative to Veh-treated group (**Figure 4H**).

**Figure 4.**
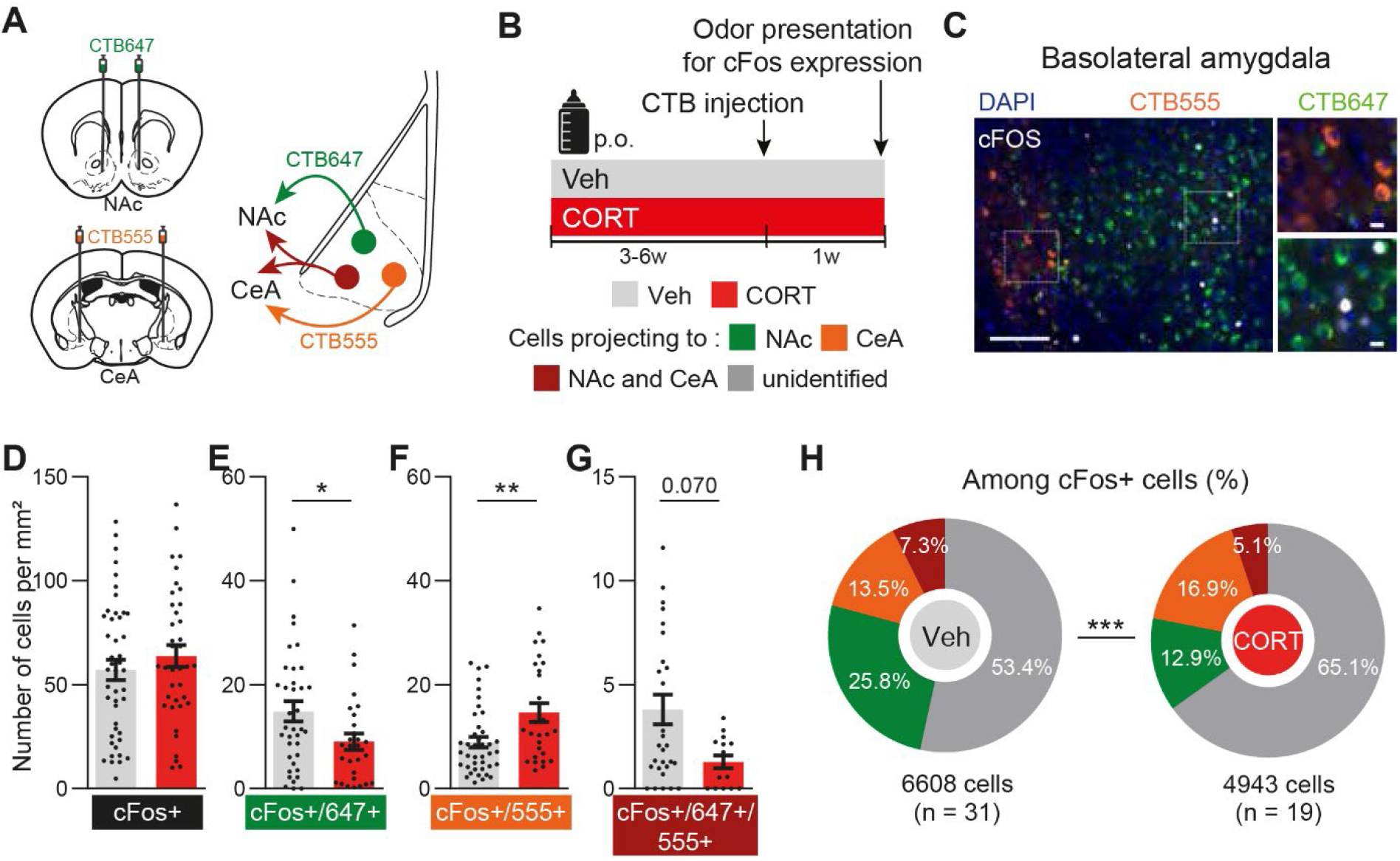
Altered BLA circuits activity after chronic CORT administration. (**A-B**) Retrograde fluorescent CTB647 and CTB555 dyes were injected in the NAc (green) and CeA (orange) respectively. Odors were presented to trigger the immediate-early gene cFos expression in CORT (red) and Veh (grey) mice. (**C**) Representative image of BLA cFos expression colocalized with CTB647 and/or CTB555. Scale bars, 100 μm (left) and 20 μm (right). (**D-H**) Quantification of cFos+ (**D**), cFos+/CTB647+ **E**), cFos+/CTB555+ (**F**) and cFos+/CTB647+/CTB555+ cell number (**G**) in the BLA. BLA cFos+ cell density was similar between groups (**D**, t(81) = 0.93, p = 0.356, n = 38-45). BLA cFos+/CTB647+ cell density decreased in CORT mice (**E**, U = 356, *p = 0.032, n = 28-37), whereas cFos+/CTB555+ cell density increased (**F**, U = 328, **p = 0.006, n = 27-40). BLA cFos+/CTB647+/CTB555+ cell density tended to decrease in CORT mice (**G**, U = 144.5, p = 0.070, n = 15-29) (**H**) Distribution of CTB647 and/or CTB555 colocalization among the total number of cFos+ cells in the BLA (χ^2^(3) = 340.0, ***p < 0.001). Bars are mean ± sem.

Previous reports demonstrated specific patterns of connection and functions associated with the basal and lateral amygdala subregions (BLA or BA and LA, respectively (26–28). However, valence coding has been inconsistently studied across these regions, analyzing either the basal part exclusively (29), or both basal and lateral parts together (13). To study specificities of each region, we performed similar analyses in the LA showing some variations compared to (basal) BLA data. Importantly, no difference were found on both LA-to-NAc and LA-to-CeA neuronal activation between Veh-treated controls and CORT-treated mice while the LA-to-NAc-and-CeA cells were more recruited in CORT-treated mice (**Figure S6** and **Table S2)**.

Although inconsistent throughtout studies, topographical gradients of cells have been suggested to distinguish their preferential positive or negative encoding role (30,29). When analyzing the spatial coordinates of cFos+ and cFos+/CTB+ cells on antero-posterior, medio-lateral and dorso-ventral axes, very few differences could be observed in the topography of these cells between CORT-treated and Veh-treated groups (**Figure S7** and **Table S3**).

In summary, chronic CORT administration induced decreased recruitment of BLA-to-NAc neurons and increased recruitment of BLA-to-CeA neurons, circuits already implicated in opposite valence encoding.

### CORT treatment induces specific alterations in presynaptic connectivity of BLA projecting neurons

To understand how depressive-like states imprinted durable anatomical changes in brain circuits, we investigated presynaptic connectivity of the BLA projecting neurons in Veh and CORT-treated mice, using targeted retrograde rabies virus-based monosynaptic tracing. In this approach, we restricted the rabies infection only to BLA-to-NAc or BLA-to-CeA neurons with retrograde Cre-expressing virus injected either in the NAc or the CeA, together with a conditional Cre-dependent ‘helper’ virus to express the avian tumor virus receptor A (TVA) and the rabies glycoprotein injected in the BLA (**Figure 5A, B**). Injected animals were treated either with CORT or Veh (**Figure 5C** and **Figure S8A-D**). Four weeks later, we injected a glycoprotein-deleted EnvA-pseudotyped mCherry-expressing rabies virus [(EnvA)SAD-ΔG-mCherry] that only infected neurons expressing TVA receptors and spread retrogradely only from cells expressing the glycoprotein, meaning specifically in presynaptic inputs of BLA-to-NAc or BLA-to-CeA neurons. Six days post-injection, we examined sections of the entire brain to quantify the location and the relative number of inputs neurons (mCherry+) (**Figure S8E**). Labelled brain regions directly projecting onto BLA-to-NAc and BLA-to-CeA neurons were similar to those previously reported (31,32) (**Figure S8F,G**). Comparing CORT with Veh-treated mice, we established a brain map of differential contributions of input regions, revealing which brain circuits exhibited distinct connectivity patterns after depressive state induction (**Figure 5D** and **Figure S8F,G**). We observed that the thalamus region (TH) and the somatomotor areas (MO) contained fewer inputs neurons to the BLA-to-NAc neurons in the CORT-treated mice (**Figure 5D top)**. Besides, we found more cells contacting BLA-to-CeA projecting neurons in the BLA circuit but a reduced the number in the pallidum (PAL; **Figure 5D bottom**).

**Figure 5.**
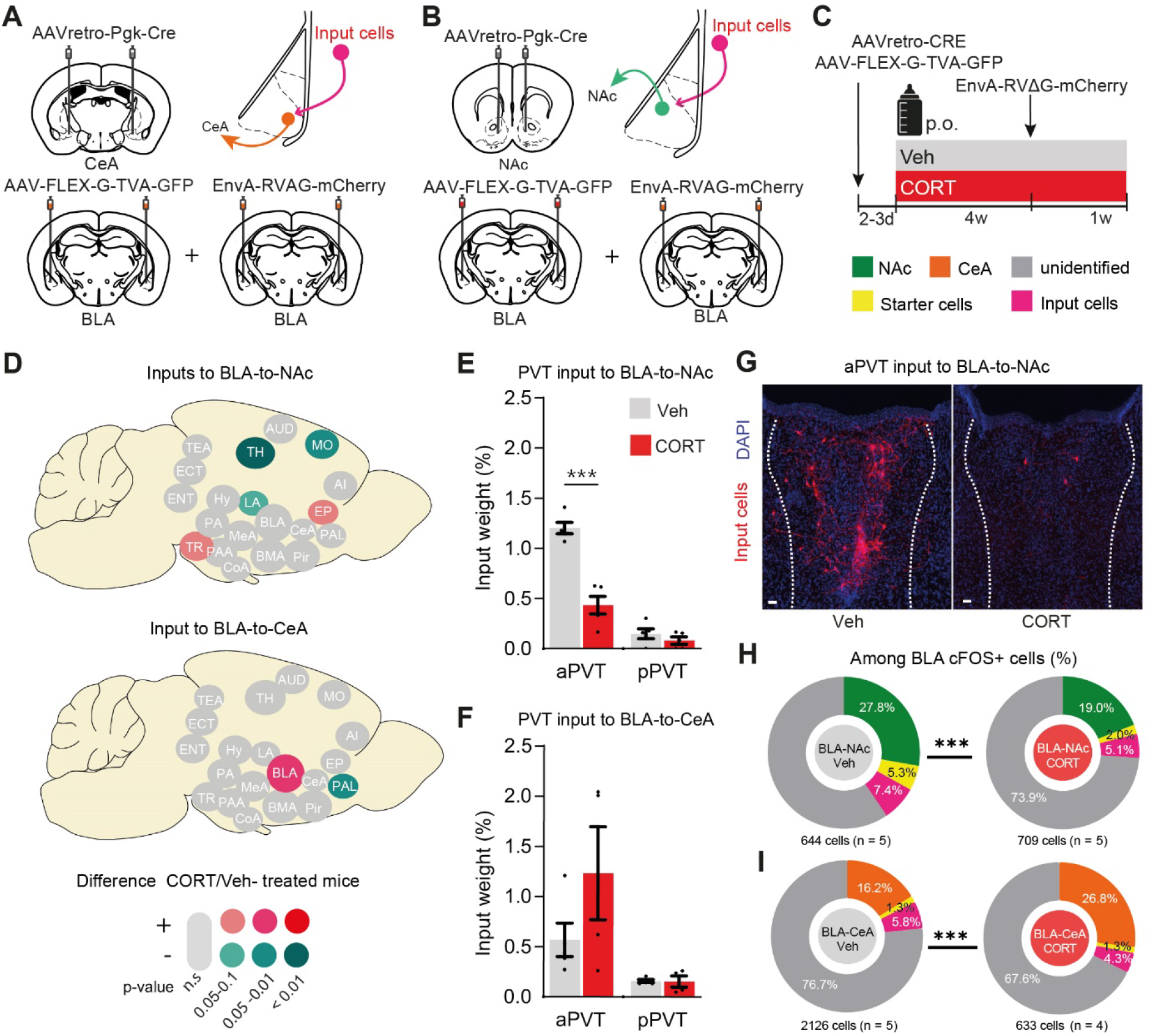
CORT treatment changes synaptic connectivity of BLA neurons. (**A**) AAVr-Pgk-Cre in the NAc and AAV-hSyn-Flex-G-TVA-GFP and EnvA-RVAG-mCherryin the BLA were injected to trace monosynaptic input to BLA-to-CeA projecting cells (in orange, n = 9). (**B**) Same than A except that AAVr-Pgk-Cre were injected in the CeA to labelled BLA-to-NAc cells (in green, n = 10). (**C**) Mice received vehicle (grey, n = 10) or chronic CORT (red, n = 9) to model depression. (**D**) Map of the differences in inputs arriving to BLA-to-NAc (*top*) and BLA-to-CeA (*bottom*) projecting neurons in CORT-treated compared to Vehicle group. Only regions showing a tendency or significant difference are colour-coded (n=4-5 per group; for full data see Figure S8F,D). (**E, F**) Percentage of presynaptic neurons to BLA-to-NAc (**E**) and BLA-to-CeA (**F**) arriving to subregions of PVT in both CORT and Vehicle groups (relative to the total). (**G**) Trans-synaptically labelled neurons in aPVT contacting BLA-to-NAc cells. (**H-I**) Distribution of BLA-to-NAc (**H**, in green) or BLA-to-CeA (**I**, in orange), starters cells (in yellow), inputs cells (in pink) and unidentifed cells (in gray) among the total number of cFos+ cells in the BLA (χ^2^(3) = 340.0, ***p < 0.001). MO: somatomotor areas; AUD: auditory areas; AI: agranular insular area; TEA: temporal association areas; ECT: ectorhinal area; PIR: piriform area; COA: cortical amygdalar area; PAA: piriform-amygdalar area; TR: postpiriform transition area; LA: lateral amygdala; BLA: basolateral amygdala; BMA: basomedial amygdala; PA: posterior amygdala; EP: endopiriform nucleus; ENT: retrohippocampal region; MEA: medial amygdala; CEA: Central amygdala; PAL: pallidum; TH: thalamus; HY hypothalamus. Bars are mean ± sem. Scale bar = 20μm.

Remarkably, more that 60% of the connections arriving from the TH in CORT-treated mice originated in the paraventricular nucleus of the thalamus (PVT) (BLA-to-NAc: 67,19 ± 3,98; BLA-to-CeA: 64,11 ± 10,58, n=4-5). CORT administration specifically reduced the number of neuronal inputs to the BLA-to-NAc neurons from the anterior PVT (aPVT) without changes from the posterior part (pPVT; **Figure 5E,G**). Interestingly, not significant difference were observed for inputs to BLA-to-CeA neurons (**Figure 5F,G**).

We also quantified in this experiment the activity of BLA specific circuits in Veh and CORT-treated mice, replicating with a different method the observation of reduced activation of BLA-to-NAc neurons in CORT-treated mice, while stronger activity of BLA-to-CeA neurons (**Figure 5H, I**, see **Figure 4H**). Overall, differential activation of BLA-to-NAc and BLA-to-CeA neurons after chronic CORT administration matches with altered presynaptic connectivity from brain regions implicated in valence assignment regulation (14).

### Chemogenetic BLA-to-CeA activation in control mice

To know if BLA circuits alterations had a causal role in the CORT-treated mice phenotype, we first attempted to elicit anxiety- and depressive-like behaviors as well as negative olfactory valence bias by chemogenetically activating BLA-to-CeA neurons in control mice. We injected retrograde AAVr-Pgk-Cre viral vector in the CeA and AAV-hSyn-DIO-hM3Dq-mCherry (or AVV-hSyn-DIO-mCherry) in the BLA to express hM3Dq, an activator designer receptor exclusively activated by designer drugs (DREADD), coupled with mCherry protein (or mCherry alone) specifically into BLA-to-CeA neurons (**Figure 6A**). Intraperitoneal (i.p.) clozapine-n-oxide (CNO) injection activated the cells (**Figure 6B**), as confirmed by increased cFos expression (**Figure S9Q-S**). Remarkably, chemogenetic BLA-to-CeA activation was not sufficient to trigger anxiety-like phenotypes in the OF and LDb, nor depressive-like phenotypes in the ST and TST (**Figure S9C-M**), as reflected by the unchanged global emotionality score (**Figure 6C**). Consistenly, it also failed to induce negative olfactory valence bias in the olfactory preference test (**Figure 6D,E** and **Figure S9N-P**).

**Figure 6.**
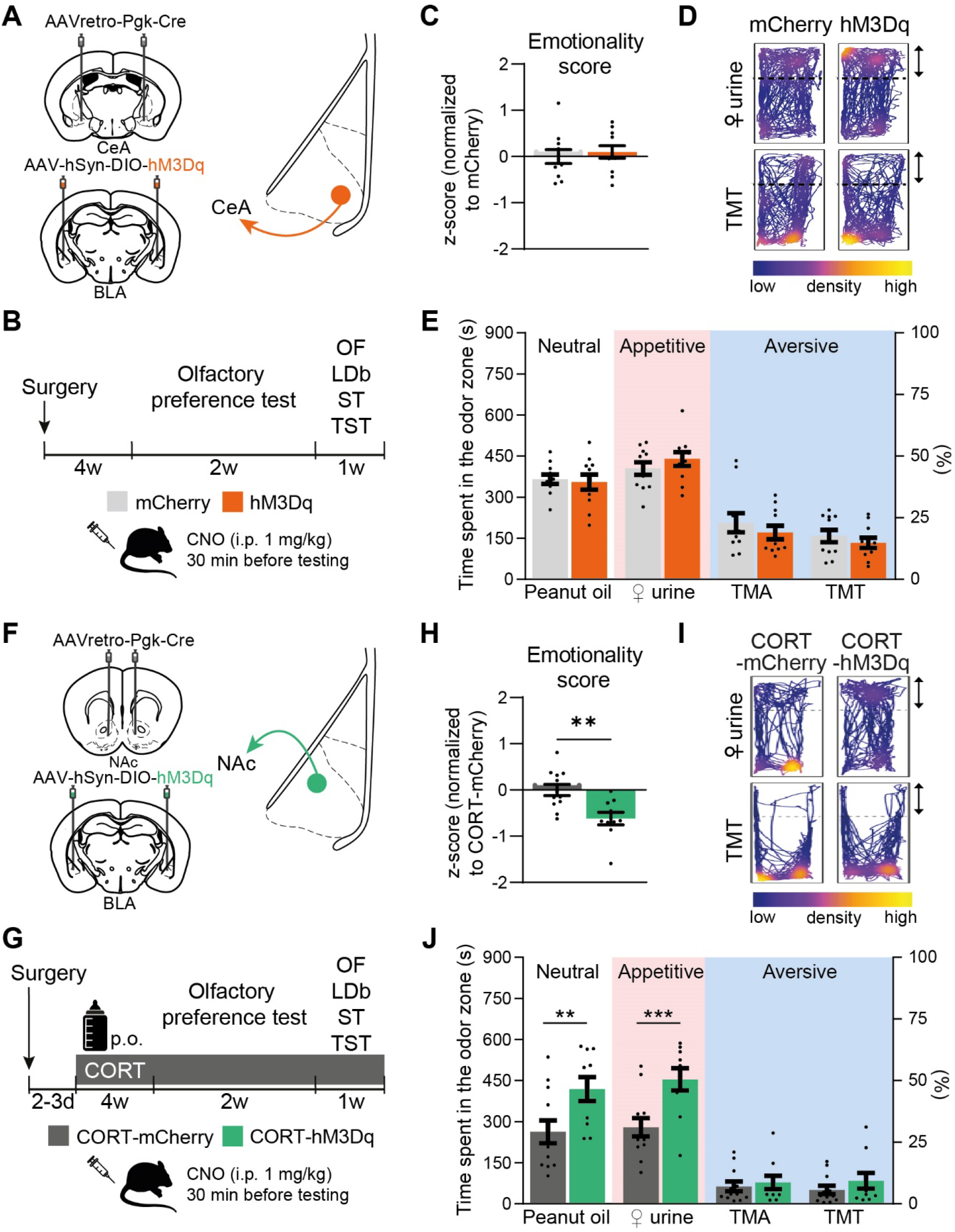
Chemogenetic activation of BLA circuits distinctly impact CORT phenotype. **(A)** To mimic chronic CORT phenotype were injected AAVr-Pgk-Cre in the CeA and AAV-hSyn-DIO-hM3Dq-mCherry (orange, n = 9, or AAV-hSyn-DIO-mCherry for the controls, grey, n = 11) in the BLA to activate BLA-to-CeA cells (**B, C**) CNO intra-peritoneal (i.p.) injection in the hM3Dq mice had no effect on global emotionality score (t(20) = 0.49, p = 0.626).). (**D**) Representative mouse tracks in the olfactory preference test. (**E**) Chemogenetic activation of BLA-to-CeA cells did not modify olfactory valence compared to mCherry controls (Group: F(1,18) = 0.10, p = 0.752; Odor: F(3,54) = 75.02, p < 0.001; Interaction: F(3,54) = 1.16, p = 0.336). (**F**) AAVr-Pgk-Cre in the NAc and AAV-hSyn-DIO-hM3Dq-mCherry (green, or AAV-hSyn-DIO-mCherry for the controls, grey) in the BLA were injected to activate BLA-to-NAc cells. (**G)** All mice were treated with CORT. (**G, H**) CNO intra-peritoneal (i.p.) injection in the hM3Dq mice reduce the global emotionality score (t(20) = 3.40). (**I**) Representative mouse tracks in the olfactory preference test. (**J**) CORT-hM3Dq mice explored more peanut oil and ♀ urine relative to CORT-mCherry controls, but not aversive odors (Group: F(1.20) = 6.56, p = 0.019; Odor: F(3,60) = 11.2, p < 0.001; Interaction: F(3,60) = 7.31, p < 0.001). **p < 0.01, ***p < 0.001. Bars are mean ± sem.

### Chemogenetic BLA-to-NAc activation in CORT mice

Secondly, we chemogenetically activated BLA-to-NAc neurons in CORT-treated mice in order to alleviate anxiety- and depressive-like phenotype, as well as the negative olfactory valence bias. We injected retrograde AAVr-Pgk-Cre viral vector in the NAc and AAV-hSyn-DIO-hM3Dq-mCherry (or AVV-hSyn-DIO-mCherry) in the BLA, to express hM3Dq coupled with mCherry protein (or mCherry alone) specifically into BLA-to-NAc neurons (**Figure 6F**). We administered CORT to all animals for four weeks and then performed behavioral testing after CNO i.p. injection (**Figure 6G**). Chemogenetic BLA-to-NAc neuronal activation, confirmed by cFOS staining (**Figure S10S-U**), was sufficient to improve anxiety-like behavior in CORT-hM3Dq mice compared to CORT-mCherry control mice in the OF but not in the LDb (**Figure S10D-H**). It had no effect on the depressive-like phenotype measured by the ST, but had antidepressant effect on the TST (**Figure S10I-N**). Overall, activating BLA-to-NAc neurons improved the global emotionality score (**Figure 6H**). Strikingly, BLA-to-NAc activation also increased exploration of peanut oil and ♀ urine in CORT-hM3Dq mice relative to CORT-mCherry controls, while leaving unchanged the aversive odors exploration (**Figure 6I,J** and **Figure S10Q,R**). It is worth noting that the time spent exploring the object during habituation also tended to be increased in CORT-hM3Dq mice (**Figure S10P**). Therefore, BLA-to-NAc neuronal activation is sufficient to improve negative olfactory bias and potentiates positive valence assignment of neutral or appetitive odors in a mouse model for depression.

## Discussion

By using a translational preference olfactory test, we demonstrate for the first time that negative emotional bias is present in a mouse model for depression similarly to BD depressed patients. Both human and mouse experiments demonstrated a shift towards more negative valence assignment of both attractive and aversive odors during depressive states. In mice, opposite alterations in BLA-to-NAc and BLA-to-CeA neurons activity correlated with this bias. Furthermore, activation of BLA-to-NAc circuit was sufficient to reverse at least partially the negative emotional bias induced by CORT treatment along with the depressive-like phenotype (**Figure S11**). Similar phenotype alleviation was observed after antidepressant administration. Additionally, we showed that CORT treatment induces specific alterations in presynaptic connectivity of BLA projecting neurons. For instance, CORT-treated mice received less neuronal inputs from the aPVT on BLA-to-NAc neurons but not significant difference in BLA-to-CeA neurons.

### Emotional bias: A common hallmark in mouse model for depression and depressed patients

Pre-clinical behavioral tests exploring valence assignment are restricted to evaluating anhedonia through sucrose preference or self-administration of psychostimulant drugs. Here, we developed a translational assay in mice assessing innate attractive and aversive odor responses, independently of any learning. We found that chronic CORT treatment induced negative shift of valence assignment on both positive (that became less attractive) and negative (that became more aversive) odors without effect on neutral odor. We observed similar bias in BD depression, such that patients rated less odors as pleasant and more odors as unpleasant than both control subjects and recovered BD patients. This negative bias correlated with depression severity. Comparably with mice, no difference was found in the valence assignment of neutral odors, suggesting bias specific towards salient emotion-triggering stimuli. Interestingly, this emotional bias regresses after recovery of the depressive episode in patients (i.e. in euthymic BD patients) and regression of depressive-like phenotype in mice responsive to FLX suggesting a major link between depressive states and the expression of negative emotional bias.

Extracting the involvement of stimulus intensity processing in our data is still elusive. Previous results have shown that chronic CORT administration induces deficits in olfactory acuity, fine discrimination of mixed odorants and olfactory memory (23). In our study, we only analyzed innate responses to individual highly concentrated odors. Furthermore, the increased response to aversive odor in CORT-treated animals reinforces the assumption that mice effectively detected these stimuli.

In human, previous data have shown that emotional bias assessed with olfactory test is present in patients suffering from unipolar depressive episode (5,33–36). One study found modifications in valence assignment in BD depressed patients, but only for pleasant odors (34). Here, we extend these data, showing that the negative emotional bias affects both positive and negative odors. Our results and previous literature indicate that negative valence attribution is a core dimension of depression whatever the type of mental disorder. Of note, we observed similar olfactory performances (detection, discrimination and identification) of BD depressive patients compared to control subjects and euthymic patients, consistently with the only comparable study (34).

Emotional biases are usually assessed in humans with visual tests such as emotion face recognition or valence assessment of pictures (37). As not doable in mice, it seemed important to verify that olfactory emotional biases were as relevant in humans as in mice. Consistently, literature on valence coding shows shared neuronal circuits for stimuli from all sensory modalities (38).

### Alteration of BLA circuits after CORT administration

To investigate the mechanisms underlying negative emotional bias observed in mice with depressive-like phenotype, we explored circuits involved in valence assignment, particularly BLA circuits. We found that BLA-to-NAc neurons expressed less cFos, an immunohistological marker for neuronal activity, in CORT-treated animals compared to the Veh-treated control group, independently of the odor stimuli used to stimulate neurons. Conversely, the density of cFos+ BLA-to-CeA neurons was higher in depressed-like mice. Some differences were found in BLA *vs* LA projecting neurons suggesting both regions could be differently affected in depression. Overall, our data underpin our hypothesis that specific BLA circuits alterations may support negative emotional bias in a mouse model for depression and concur with previous studies showing that BLA-to-NAc neurons preferentially encode positive valence stimuli whereas BLA-to-CeA preferentially encode negative valence stimuli (12–14), but see also (16,39).

What are the mechanisms behind BLA circuit dysregulation in depressive states? BLA-to-NAc and BLA-to CeA population neurons are highly interconnected and they mutually control their activity (30,40). Moreover, although less abundant, local interneurons can tightly tune BLA circuits functioning (41). Chronic stress and increased peripheral cortisol/corticosterone levels might disrupt molecular and cellular neuroplasticity in the BLA affecting both principal neurons and local interneurons. Glucocorticoid receptors are located in all portions of the amygdala (42,43). Consistently, chronic stress dramatically impacts BLA principal neurons both at morphological and physiological levels (27). Besides the BLA, CORT treatment could affect other brain regions indirectly affecting these circuits. For instance, prolonged corticosteroid stimulation leads to atrophy of apical dendrites in the hippocampus (27,44–46) and is associated with elimination of postsynaptic dendritic spines in medial prefrontal cortex projection neurons (47).

Using rabies virus-based retrograde tracing, we observed modifications of presynaptic inputs to both BLA-to-NAc and BLA-to-CeA neurons following CORT administration. Notably, aPVT, but not pPVT, inputs to BLA-to-NAc neurons were reduced in CORT-treated mice. These results are particularly interesting in the context of a recent study demonstrating the fundamental role of PVT-to-BLA afferents for valence assignment, controlled through neurotensin secretion rate (14). Importantly, CORT receptors are also express in PVT neurons affecting their function (48). Our results suggest direct role of inputs from the PVT to BLA-to-NAc-projecting neurons in depressive states and associated negative emotional bias. Indeed, reduced afferents from PVT to BLA-to-NAc neurons could be responsible for their reduced activity rates. Further experiments will be required to identify the precise causal mechanisms behind the alteration in the different BLA circuits activity.

### Activation of BLA-to-NAc pathway partially restores negative bias

We observed increased activity of BLA-to-CeA neurons in CORT-treated mice suggesting that depressive-like phenotype and negative emotional bias could be mimicked by over-stimulation of this circuit. However, chemogenetically stimulating the BLA-to-CeA pathway in control animals induced neither anxiety- or depressive-like phenotypes, nor a negative olfactory bias. These data confirm that the anxiety/depressive-like phenotype is highly related to emotional bias.

In contrast, activating the BLA-to-NAc circuit reduced the anxiety/depressive-like phenotype in CORT-treated mice and increased the attractiveness of both neutral and positive odors. Importantly, BLA-to-NAc neurons are primarily suggested to be involved in reward processing and positive valence assignment, consistently with the effect we observed. However, activating the BLA-to-NAc neurons did not modify the aversion towards negative odors, indicating that other neuronal pathways could be required to restore the bias regarding aversive odors. Negative stimuli predicting danger might rely on multiple evolutionally selected and redundant mechanisms more difficult to hijack. Present data demonstrate that BLA-to-NAc activation is able to improve positive valence assignment, providing a mechanism to alleviate anxiety/depressive-like phenotype in CORT-treated mice (6). This neuronal activity manipulation mimicked the action of FLX in responsive mice. Together with previous studies (49,50), our results suggest BLA-to-NAc neurons might represent common pathway for antidepressant action.

In summary, we show for the first time a negative bias in valence assignment in a mouse model for depression recapitulating the emotional bias found in depressed patients. This bias is related to plasticity in BLA circuits known to be involved in valence assignment. Futures steps will require to explore the effect of various classes of antidepressants on negative emotional bias and BLA circuits activity. Identifying a common behavioral and neuronal mechanism of action for different antidepressant should be of great help for improving therapeutic management of depression. If reversibility of emotional negative bias is mandatory to expect antidepressant action, new molecules should be assessed in pre-clinical studies using olfactory valence evaluation, complementing the gold standard but challenged forced swim and tail suspension tests (51).

## Acknowledgements

We thank Denis David for advices on the corticosterone-induced model for depression, the Institut Pasteur Central animal facility and especially Noémi Dominique for her assistance with behavioral experiments, Corentin Guérinot for his advices in the PCA analyses and the Institut Pasteur Image analysis hub for help in developing the Icy protocol. We are grateful to Karl-Klaus Conzelmann (Max Von Pettenkofer Institute Virology and Gene Center, Medical Faculty, Ludwig-Maximilians-University Munich, Germany) for the generous gift of (EnvA)SAD-ΔG-mCherry virus. This work was supported by Agence Nationale de la Recherche Grants ANR-15-NEUC-0004-02 “Circuit-OPL”, Laboratory for Excellence Programme “Revive” Grant ANR-10-LABX-73, ANR-AAPG2021 “EMOKET”, the Life Insurance Company AG2R-La Mondiale, Labex BioPsy and the Fondation Deniker.

## Author contributions

MB: Conceptualization, Methodology, Investigation, Formal analysis, Visualization, Writing – Original Draft, Writing – Review and Editing; CHBD, AB, EV, MM: Investigation, Formal analysis, Visualization; PML, JH: Writing – Review and Editing, Supervision, Funding acquisition; MA: Conceptualization, Methodology, Investigation, Formal analysis, Visualization, Writing – Original Draft, Writing – Review and Editing, Supervision; CH: Conceptualization, Methodology, Writing – Original Draft, Writing – Review and Editing, Supervision, Funding acquisition.

