## Supplementary information for "Basolateral amygdala circuits supporting negative emotional bias in a mouse model for depression"

### Supplemental information

#### Methods and Materials

##### Human subjects

Research involving human research participants have been performed in accordance with the Declaration of Helsinki. The Human Research Ethics Committee, CPP-Ile de France IV (2015/44), approved the study and all participants gave their informed consent.

The study was conducted in a tertiary-care psychiatric hospital located in Créteil, France. Consecutive sampling over a 36-month period was used to recruit patients. Forty-eight patients were included in our study, aged from 18–65 years. Potential participants were independently diagnosed with a current bipolar disorder diagnosis by two board-certified psychiatrists with the aid of a structured interview based on DSM-IV criteria (1). Among the recruited patients, twenty-five subjects presented an euthymic state and twenty-three a depressive or mixed state at the moment of the evaluation. Depression severity was measured by MADRS scale. Mania severity was measured by YMRS scale.

The same two psychiatrists classified the BP individuals as BP I (n = 29) or BP II (n = 18) according to the DSM-IV criteria. A control group (n = 11 subjects) was recruited through word-of-mouth from local community.

##### Olfactory assessment

The Sniffin' sticks test (Burghardt®, Wedel, Germany) comprises 3 subtests, resulting in 3 scores: detection threshold, discrimination and identification scores (2,3).

The detection threshold score was assessed using 16 dilutions prepared from a 4% n-butanol solution (dilution ratio 1:2). Three pens (two containing the solvent and the third the odorant) were presented in a randomized order using a single staircase of increasing concentration [16 (lower concentration) to 1 (higher concentration)]. Subjects had to identify the odor-containing pen. Reversal of the staircase was triggered when the odorant was correctly identified in two successive trials. The detection threshold score was defined as the mean of the last four of seven staircase reversals, scores ranging from 1 to 16 (the higher, the better). Subjects were blindfolded during this test.

To assess the discrimination score, triplets of pens were presented in a randomized order (two containing the same and one a different odorant). Subjects had to determine which of three pens smelled differently, scores ranging from 1 to 16. Subjects were blindfolded during this test as well.

The identification score was assessed for 16 common odors (orange, leather, cinnamon, peppermint, banana, lemon, liquorice, turpentine, garlic, coffee, apple, cloves, pineapple, rose, anise and fish). Using a multiple-choice task, the identification index of individual odors was performed from lists of four descriptors each, scores ranging from 1 to 16. Moreover, during the Sniffin' sticks identification task, patients and control subjects were asked for the pleasantness of the 16 selected odors smelling, as it was previously published (4). The number of odors rated as 'pleasant', 'unpleasant' or 'neutral' range from 0 to 16 and the sum of these scores was equal to 16.

These obtained results could not be explain by tobacco use that do not differ between groups (Two-way RM ANOVA; Smoking status: $F(1,57) = 0.00$, $p > 0.999$, Valence: $F(2,114) = 33.96$, $p < 0.001$, Interaction: $F(2,114) = 2.27$, $p = 0.108$).

**Animals**

All animal care and experimental procedures followed national and European (2010/63/EU) guidelines and were approved by the French Ministry of Research (APAFiS: #16380-2018080217358599_v1). C57BL/6N male mice (7–8 weeks old) purchased from Taconic Farms (Denmark) were used for all behavioral tests and immunohistochemistry studies (n = 181). Once arrived in the animal facility, they were given at least one week to accommodate. Mice were socially housed, 4-6 per cage, and maintained under standard housing conditions (23 ± 1 °C; humidity 40%) on a 14/10 h light/dark cycle (lights were on from 7 AM to 9 PM every day) with food and water *ad libitum*, except during behavioral experiments and odor exposures. All behavioral tests were conducted during the period of light (10 AM-7 PM). We used the minimum number of animals, estimated from our previous knowledge in performing the same type of experiments.

**Drugs**

**Chronic corticosterone administration depression model**

Corticosterone (CORT) purchased from Sigma-Aldrich (France) was sonicated for 2 h in a vehicle made of 10% (2-hydroxypropyl)-beta-cyclodextrin (β-CD; Sigma-Aldrich, France) in water. After complete dissolution, the solution was added to the appropriate amount of water to reach the final concentration of 35 µg/ml CORT and 0.45% β-CD. CORT (35 µg/ml, equivalent to about 5 mg/kg/day) or vehicle (0.45% β-CD) was available *ad libitum* in the drinking water in bottles wrapped with aluminum to protect it from light. For fluoxetine experiments, after four weeks of CORT treatment, fluoxetine hydrochloride (equivalent to about 18 mg/kg/day; Sigma-Aldrich) was administered during three weeks for antidepressant-like effects together with CORT administration.

All the bottles were changed every 3-4 days in order to prevent any possible degradation, as previously described (5). Animal were weighted twice a week to verify the increase in body mass already described in this model (5).

**Clozapine n-oxide**

Clozapine n-oxide (CNO) dihydrochloride (Tocris, France) was dissolved in sterile saline (NaCl 0.9%) and adjusted to a final concentration of 0.2 mg/ml. On behavioral testing days, mice received intra-peritoneal (i.p.) injection at a concentration of 1 mg/kg, 30 min before testing. Fresh CNO solution was prepared every 1-3 days.

**Stereotaxic surgery**

Buprenorphine (0.05 mg/kg sub-cutaneous (s.c.) injection) was administered 30 minutes before the surgery. Mice were anesthetized with a ketamine/xylazine mix (150 mg/kg and 5 mg/kg respectively, i.p. injection) and placed in a stereotaxic frame, with their body temperatures maintained using a heating pad. The cranial skin was shaved and disinfected with povidone iodine, and craniotomies were performed above the injection sites. Dropplets of lidocaine were administered on the cranial skin before aperture and suture. At the end of the surgery, meloxicam (1 mg/kg s.c. injection) was administered to help recovering.

**CTB injection**

In order to label the neurons of the Basolateral Amygdala (BLA) projecting to the Nucleus Accumbens (NAc) or the Central Amygdala (CeA), the retrograde tracer Cholera Toxin Subunit B conjugated with Alexa Fluor 555 or 674 (CTB 555 or CTB 647, 1 mg/ml, Invitrogen, USA), was bilaterally injected with a glass micropipette connected to a Nanoject III microinjector (Drummond Scientific) at a rate of 5 nl/s in the NAc (AP : +1.4 mm ; ML : ±0.87 mm ; DV : −4.35 mm, 300 nl) or in the CeA (AP : -0.8 mm, ML : ±2.35mm, DV : -4.35 mm, 100 nl, targeting the centromedial part). Distances were measured from theoretical bregma and brain surface. The needle was left in place one minute before each injection and 2-5 min after, and then slowly withdrawn. Allen Brain atlas was use as reference to report injection site in each animal (**Figure S4**). Brain hemispheres where the injection site was outside the analyzed regions were discarded for the analysis (NAc: Hemispheres, n = 50/166, Mice, n = 17/83; CeA: Hemispheres, n = 71/166, Mice, n = 17/83).

**Viral vector injection**

For labelling the presynaptic inputs to BLA-to-NAc and BLA-to-CeA cells, AAV-FLEX-G-TVA-GFP was injected in the BLA (100 nl), and AAVretro-CRE either in the NAc (100 nl) or in the CeA (50 nl) using the previous coordinates. After the four weeks of CORT treatment, mice were injected with EnvA-RVΔG-mCherry in the BLA (150 nl) and sacrificed one week later.

AAV5-hSyn-DIO-hM3Dq-mCherry, AAV5-hSyn-DIO-mCherry and AAVretro-PGK-Cre recombinant adeno-associated virus were used to manipulate the BLA neuronal activity. The viral vectors were bilaterally injected into the BLA (AP: −1.75 mm, ML: ±3.15 mm, DV: −4.25 mm, 100-150 nl) and the NAc (same coordinates as mentioned above, 100-150 nl) or the CeA (same coordinates as mentioned above, 50 nl) as previously mentioned. Animals whom both hemispheres were injected outside the analyzed regions were discarded for the analysis (BLA-to-NAc: n = 2/24; BLA-to-CeA: n = 2/22).

All AAV were purchased from Addgene (USA) ) and EnvA-RVΔG-mCherry was generated by Karl-Klaus Conzelmann lab (University of Munich, Germany).

**Behavioral assessment**

A battery of behavioral tests was used to assess anxiety and depression-like phenotypes, and olfactory preference. Before starting behavioral testing, mice were handled ~30 s at least twice a day for 3 days to habituate to the experimenter.

**Open-field**. Animals were placed in grey Plexiglas containers (43 x 43 cm2) and their behavior was recorded by a video camera during 20 min. A tracking system (Noldus Ethovision 3.0, Netherlands) was used to map center and periphery zones and to calculate the time spent and distance moved in each zone. The time spent in the center, the number of entries in the center and the total distance traveled were calculated as measures of anxiety behavior and ambulatory activity respectively. The arenas were cleaned with water between each trial.

**Light and dark box**. A two-compartment box containing a dark chamber (black walls with upper lid) and a light chamber (~300 lux, white Plexiglas walls, no upper lid) was used. The chambers were connected by a 10 × 10 cm door in the middle of the wall. Animals were placed in one corner of the light chamber facing the wall and were allowed to freely explore for 6 min. The Noldus Ethovision 3.0 tracking system was used to record behavior. The number of entries and time spent in the light chamber were estimated as measures of anxiety. Between each trial, the light/dark compartments were cleaned with water.

**Splash test**. The test consisted in squirting ~ 200 µl of a 10% sucrose solution on the dorsal coat of the mouse and placing it in its home cage without cage mates. The test was performed in a quiet, dimly lit room (~30 lux) and video-recorded. Grooming latency, frequency, and duration were assessed during 6 min by blind experimenter, as measures of self-neglect and depressive-like behaviors.

**Tail suspension test**. Mice were suspended at approximately one-third from the end of the tail, using regular tape, to a metal rod about 30 cm from the table, for 6 min. Upon viewing of the video recordings blindly to the treatment, the latency to immobility, the number of immobility episodes and the total time spent in an immobile posture was measured. Longer periods of immobility are associated with depressive-like states.

**Coat state test**. The total coat state score resulted from the sum of the score of five different body parts:  neck, dorsal/ventral coat, tail, forepaws and hindpaws. For each body area, a score of 0 was given for a well-groomed coat and 1 for an unkempt coat (6,7). The measurements of the coat state were done by an experimenter blind to treatments.

**Olfactory preference test**. The test was adapted from Pérez-Gómez et al. (8). The test was performed in a quiet and dimly lit room (~ 40 lux), around 3 to 6 pm. Clean housing cages (17 x 32 cm) with regular bedding material were used as testing arenas, covered by transparent Plexiglas lids. Each testing arena received mice socially housed in the same cage. The first day, all the mice from the same cage were placed together to overcome neophobia, in each testing arena for habituation during 15-20 min. The second day, a petri dish (94 mm diameter) with a hooled cover was placed and adhered to one side of the arena, for defining an odor zone (one-third of the cage). For 12 consecutive days, the behavior was recorded by a video camera for 15 min and the Noldus Ethovision 3.0 system was used to track the position of the mice. The time spent in an odor zone and the locomotor activity were used as measures of olfactory valence. During the first 4 days, only a Whatman paper filter (GE Healthcare Life Sciences, USA) was placed into the petri dish, to assess the baseline exploration. Animals exploring less than 225 s the odor zone during habituation (25%) were excluded from the analyses. Then, 2 days were dedicated to each odor, placed on a paper filter, in the following order: peanut oil (pure, 400 µl), female urine (pure, 100 µl), trimethylamine (Sigma-Aldrich, 6.75% in water, 400 µl) and 2,4,5-trimethylthiazole (Sigma-Aldrich, 5% in mineral oil, 400 µl). Peanut oil and female urine are classical appetitive odorants, whereas trimethylamine and 2,4,5-trimethylthiazole are predator urine synthetic compounds, aversive at these concentrations (8–10). However, in our experimental conditions, animals were not food deprived before the test and consequently peanut oil had a neutral value and did not trigger attraction. Aversive odors were kept for the last days of the test in order to avoid any potential aversive conditioning. Repeating the measurement of each odor twice allowed to decrease the inter-subject variability of this spontaneous behavior.

**Behavioral tests z-scores**

z-scores were calculated by standardization of the values measured (substracted by the mean of the control group [Veh mice in **Figure 2**, BLA-to-CeA mCherry mice in **Figure 6C** and BLA-to-NAc CORT mCherry mice in **Figure 6H**], and then divided by the control group standard deviation) for each parameter of each behavioral test. The directionality (*i.e.* the positive or negative sign) of the z-scores was adjusted so that an increased value corresponds to more anxiety- or depressive-like phenotype (11). It included the time spent and the number of entries in center and the distance moved for the open field, the time spent and the number of entries in the light box for the light and dark box, the latency to first grooming, the number of grooming episodes and the time spent grooming for the splash test, and the latency to immobility, the number of immobility episodes and the time spent immobile for the tail suspension test. The global z-scores for each behavioral test were calculated as the mean of z-scores of each parameter measured in the test.

**Odor stimulation for cFos activation**

Exposure sessions were conducted in custom-made device that allow a specific and controlled odor stimulation avoiding odor contaminations. Animal were located in small cages (0.5 L) that could be rapidly saturated with odorant vapor (pump air flux 3.5 L/min; total saturation time ~ 9 s). The odor input was located on the top of the cage and an extraction fan was located on the bottom to evacuate the odorant vapor (fan speed, 147 L/min; total evacuation time ~ 200 ms). Mice were first habituated to the device for 1 h for 3 consecutive days. During these habituation sessions, animals were subjected the first day to deodorized air with the extraction fan operating continuously. During the second and third days of habituation, the protocol alternating the functioning of fan and pump as in the day of test was used. The day of test, mice were subjected to (1) deodorized air for 60 min to minimize basal cFos expression, (2) odorant-containing air pulses (0.3 L/min pump odor flux; 8.5% final odor dilution) for 60 min (36 pulses of 40 s each, interleaved with 60 s applications of deodorized air to avoid sensory habituation), (3) additional 2 h in the small cages with pump and fans off. The mice were then quickly perfused. Animals from Veh and CORT-treated groups were exposed with only one odor solution: mineral oil (pure, 10 ml), female urine (pure, 5 ml) or TMT (10% in mineral oil, 10 ml). For female urine, a cotton stub impregnated with 100 µl of female urine was located on the top of the cage and the bottle containing the female urine solution was heated at 37 °C to increase odor stimulation.

**Immunohistochemistry**

Mice were deeply anesthetized with a ketamine/xylazine mix (i.p., 150 mg/kg and 50 mg/kg respectively) and perfused transcardially with a solution containing 0.9% NaCl, followed by 4% paraformaldehyde in phosphate buffer, pH 7.3. Brains were removed and postfixed by incubation in the same fixative at 4°C for additional 2 hours. Brain were then cryoprotected by incubation in 30% sucrose + 0.02% azide in PBS for 24-48 h. Forty-micrometer-thick coronal brain sections were obtained using a sliding microtome (Leica SM 2010 R). Immunostaining was performed on free-floating sections. Non-specific staining was blocked by 0.2% PBS-Triton and 10% normal donkey serum (Sigma-Aldrich, Germany). Sections were incubated with 0.2% PBS-Triton, 4% bovine albumin serum (Sigma-Aldrich), 3% normal donkey serum and the following primary antibodies at 4°C : guinea pig anti-cFos (1:2000, Synaptic Systems, Germany), chicken anti-GFP (1:1000, Abcam, USA) and rabbit anti-RFP (1:4000, Rockland, USA) for 48 h. Sections were then incubated with secondary antibodies (Alexa-conjugated secondary antibodies at 1:1000, Jackson ImmunoResearch Laboratories, United Kindom and Molecular Probes, USA) at room temperature for 2 h. Fluorescent sections were stained with the nuclear dye DAPI (1:5000, Invitrogen) and then mounted using Fluoromount aqueous mounting medium (Sigma-Aldrich).

**Image acquisition**

**CTB tracing experiments.** For slices stained for cFos expression after CTB647 and CTB555 injections, tissue sections from the whole BLA (basolateral and lateral amygdala, 10-20 per mouse) were collected using an Axioscan microscope (Zeiss, 10X objective). cFos+ cells, CTB647+ and CTB555+ cells, cFos+/CTB647+ and cFos/CTB555+ double-positive cells, and cFos+/CTB647+/CTB555+ triple-positive cells in the BLA were counted automatically using Icy, an open community platform for bioimage analysis (Institut Pasteur, see Icy software protocol). Values were given as cell density (number of cells per mm²). For cFos+ cell density analysis, mean values obtained from individual mice were reported (**Figure 4, Figure S5, S6, S7**).

**Input tracing analysis**. For sections stained after EnvA-RVΔG-mCherry injection in the BLA, we imaged sections from the whole brain (about 40 sections per mouse) using an Axioscan microscope (Zeiss, 10X objective). mCherry+ cells, GFP+ cells and mCherry+/GFP+ double-positive cells were manually counted depending on their localization in the brain. To facilitate cells localization, we used the Aligning Big Brains & Atlases project (ABBA, Bioimaging and Optics Platform, Lausanne) to align each brain section to the Allen Brain atlas. It consisted in a Fiji (ImageJ 1.53f51) plugin associated with QuPath (0.2.3 version) tools. cFos+ cells were also automatically counted using QuPath in some defined brain regions, as well as mCherry+/cFos+ cells, GFP+/cFos+ cells and GFP+/mCherry+/cFos+.

**Validation of chemogenetic activation.** To verify neuronal activation of hM3Dq-expressing cells (**Figure S9 and S10**), one week after the end of behavioral experiments, animals were injected with CNO 1 mg/kg i.p. 2 h before transcardial perfusion. Immunohistochemistry for cFos and mCherry was performed as previously described. Images of slices were collected using a confocal laser-scanning microscope (LSM 700; Zeiss; 25X objective; 6-10 slices per mice). cFos+ cells and cFos+/mCherry+ double-positive cells were manually counted throughout the entire stack of optical slices (4 stacks spaced by 5 µm) by an experimenter blind to the image condition. Values were reported as total number of mCherry+ cells per field and percentage of cFos+ cells among mCherry+ cells averaged for each animal.

**Icy and R softwares**

Image analyses were performed using the Icy open source platform (http://www.icy.bioimageanalysis.org (12,13)). Regions of interest (ROI) were first drawn manually around the lateral and the basolateral amygdala. Identification of the antero-posterior coordinates of each slice was also performed manually. An Icy protocol was designed to automatically count cFos+ cells and their colocalization with CTB555 or CTB647. To do so, the "Spot Detector" plug-in was applied to each ROI (LA and BLA), creating small ROI ("spots") around each cFos+ nucleus. These spots were then dilated by 3 pixels on X and Y to include the soma of the cells. In these enlarged spots, thresholds on red (CTB555) and far red (CTB647) fluorescence intensity and standard deviation were applied to determine if the spots were CTB555+ or CTB555- and CTB647+ or CTB647-.

Parameters were set manually and were identical for all the images from one experiment replicate (n = 3 experiment replicates). Collected data sheets contained information about spot size, fluorescence intensity and standard deviation, as well as X and Y coordinates relative to the center of the ROI (LA/BLA taken together). Spatial distribution of detected cells in medio-lateral and dorso-ventral axes were calculated with respect to the center of the drawn ROI corresponding to the LA/BLA), on each coronal slice whose antero-posterior position along the LA/BLA was manually determined.

The R software (CRAN project, v3.6.3) was used to manage the data sheets generated by Icy software and produce final tables.

**Statistical analysis**

Statistical analyses were performed with GraphPad Prism v9 software (USA). Normality was assessed using the Shapiro-Wilk test. The Barlett test was then used to statistically compare the variances. Two-sided parametric or non-parametric tests were used accordingly: unpaired Student with or without Welch correction or Mann-Whitney tests, One-way ANOVA or Kruskal Wallis tests with repeated measures when suitable, Two-way ANOVA and Three-way ANOVA or Mixed-effect model when some values were missing, with repeated measures when suitable. Following post-hoc analyses were always applied with the False Rate Discovery method of Benjamini-Hochberg. Correlation analyses in **Figure 1** were performed with the Pearson or the Spearman coefficient depending on the normality of the data. Chi-square tests were performed when suitable with contingency tables. Log-rank tests were performed on survival tables.

Principal Component Analysis (PCA) was performed with R (package FactoMineR) on a dataset containing 87 animals (44 Veh-treated and 43 CORT-treated) assessed on 23 behavioral parameters (3 parameters of the open field, 2 of the light and dark box, 3 of the splash test, 3 of the tail suspension test (reported in **Figure S1**) and 12 of the olfactory preference test including for all the odors the time exploring, the mean position and the distance moved). Each behavioral parameter was z-scored as explained previously based on the Veh-treated control group of the experiment replicate (n = 3 experiment replicates). Missing values were completed using the missMDA package from R. The values were standardized so that the final mean and the standard deviation of each parameter equal 0 and 1 respectively. Six PCs were kept in order to attend > 70% of explained variance (72.6%). The contributions of each behavioral parameter to each PC were computed and summed by behavioral test. The contributions of each test to each PC are represented by the proportion of the total bar plotted in **Figure 2H**. A logistic principal component regression (PCR) on the group (Veh vs CORT) was then performed using the PCA scores of each animal for the six first PCs, and their second-level interactions. The stepAIC function from the MASS package was used to perform a stepwise algorithm in order to choose the final model based on the best Akaike information criterion (AIC), an estimator of the model quality taking into account the goodness of fit (amount of variance explained) and the simplicity of the model (number of explaining variables). The direction of the stepwise search was set on both backward and forward, meaning that removed explaining variables could be add again on each step. The final coefficient estimates and the associated p-values for the retained PCs and their second-level interactions are reported in **Figure 2I**.

All datasets were described using the mean; error bars in the figures represent standard error mean (SEM), except for **Figure S5E-G** where box-and-whiskers plots were used; boxes represent the $25^{th}$, median and $75^{th}$ percentiles while the whiskers represent the $1^{st}$ and $99^{th}$ percentiles. Differences were considered significant for $p < 0.05$.

The datasets and codes that support the findings of this study are available from the corresponding author upon reasonable request.

**Supplementary Figures and Tables**

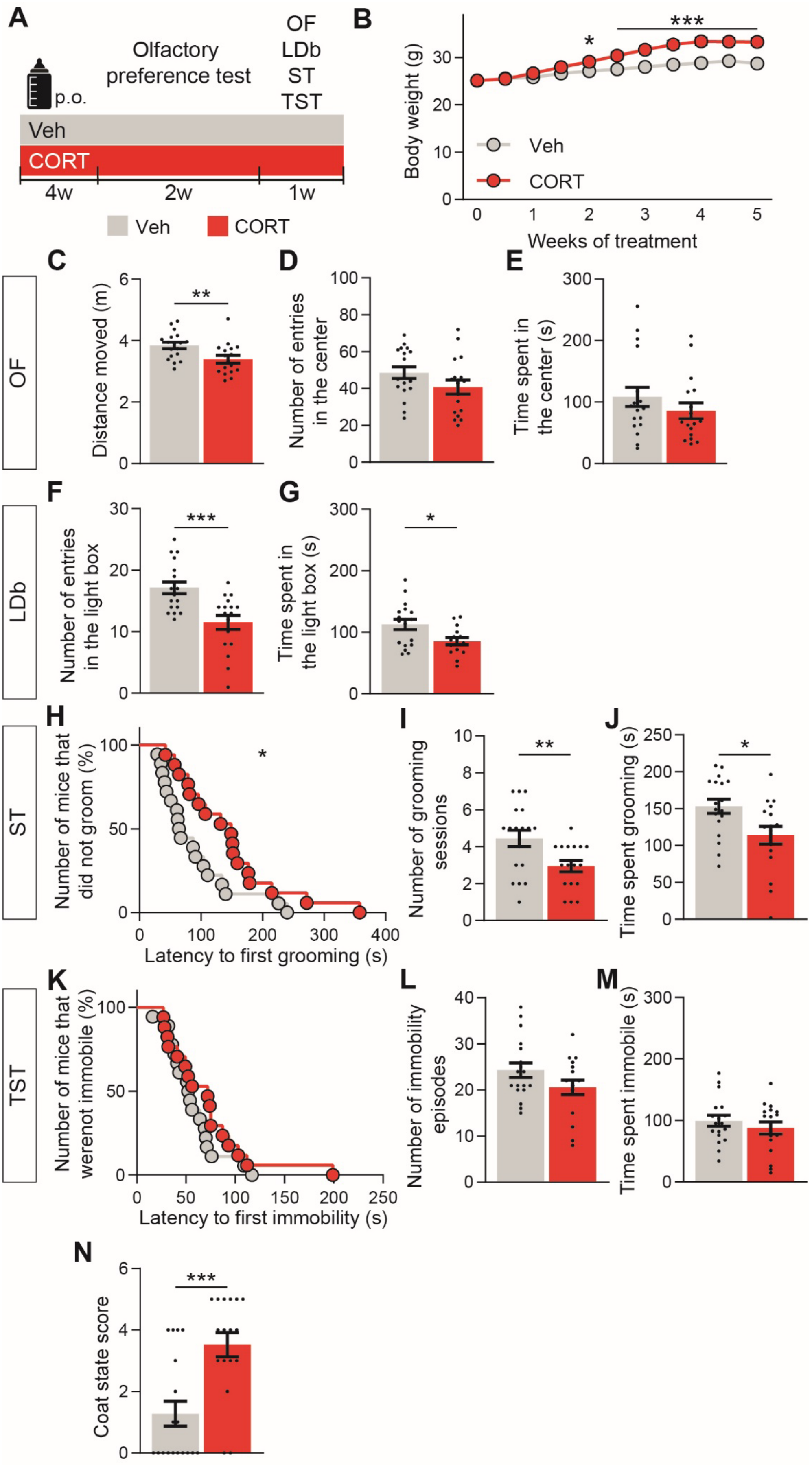
A
p.o.
Olfactory
preference test
OF
LDb
ST
TST
Veh
CORT
4w
2w
1w
Veh
CORT
B
***
*
Body weight (g)
Veh
CORT
Weeks of treatment
C
OF
Distance moved (m)
**
D
Number of entries
in the center
E
Time spent in
the center (s)
F
LDb
Number of entries
in the light box
***
G
Time spent in
the light box (s)
*
H
ST
Number of mice that
did not groom (%)
*
Latency to first grooming (s)
I
Number of grooming
sessions
**
J
Time spent grooming (s)
*
K
TST
Number of mice that
werenot immobile (%)
Latency to first immobility (s)
L
Number of immobility
episodes
M
Time spent immobile (s)
N
Coat state score
***

**Figure S1.** Chronic CORT administration triggered anxiety-like and depressive-like behaviors in mice. (**A**) Mice receive vehicle (grey, n = 18) or chronic CORT (red, n = 17) to model depression. (**B**) Chronic CORT induced more body weight gain than Veh (Group: F(1,33) = 10.61, p = 0.003; Weeks: F(10,330) = 316.3, p < 0.001; Interaction: F(10,330) = 48.10, p < 0.001). (**C-E**) CORT mice moved less in the OF compared to Veh controls (**C**, t(33) = 2.79, **p = 0.009), but did not explore differently the center (**D**, t(33) = 1.58, p = 0.123; **E**, t(33) = 1.11, p = 0.274). (**F-G**) CORT mice explore less the light box during LDb (**F**, t(33) = 3.84, ***p < 0.001; **G**: t(31) = 2.58, *p = 0.015, n = 15 CORT). (**H-J**) Grooming is reduced in CORT mice during ST, with higher latency to first grooming (**H**, $\chi^2(1)$ = 4.51, *p = 0.034) and lower grooming sessions number (**I**, t(33) = 2.76, **p = 0.009) and time spent grooming (**J**, t(33) = 2.58, *p = 0.015). (**K-M**) Neither the latency to first immobility (**K**, $\chi^2(1)$ = 1.19, p = 0.275), nor the number of immobility episodes (**L**, t(33) = 1.67, p = 0.104) nor the time spent immobile (**M**, t(33) = 0.87, p = 0.393) are different between CORT mice and Veh controls in the TST. (**N**) The CORT mice coat is deteriorated compared to Veh controls (U = 57.50, ***p < 0.001).

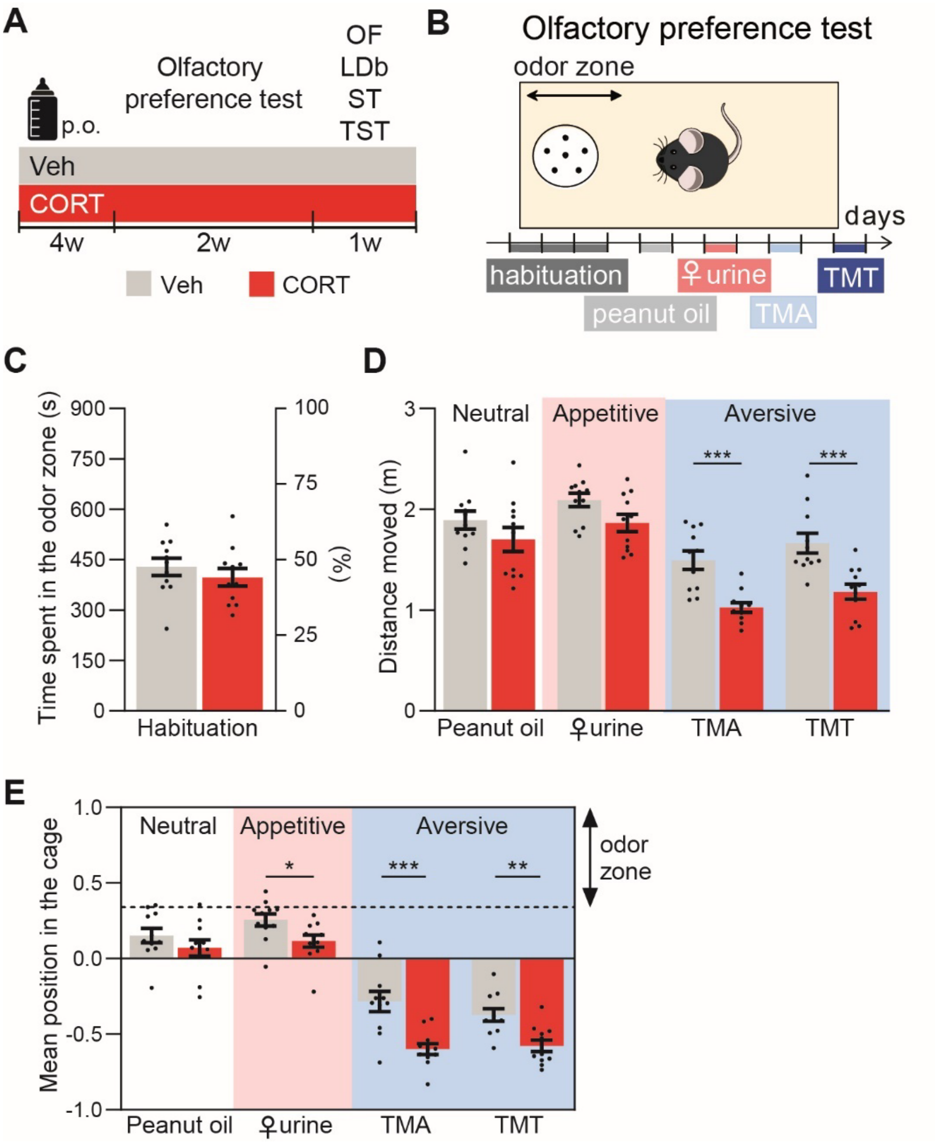


**Figure S2.** Reduced locomotor activity in CORT mice upon aversive odor presentation in the olfactory preference test. (**A**) Chronic CORT (red, n = 11) administration is used to model depression. Control mice only receive vehicle (grey, n = 11). (**B**) Scheme of the olfactory preference test protocol. (**C**) Chronic CORT administration does not alter exploration of the "object" zone habituation without odor. (**D**) Aversive odors reduce the distance moved compared to habituation (One-way RM ANOVA: F(4,40) = 18.17, p < 0.001, followed by FDR post-hoc comparisons, TMA: t(40) = 4.78, p < 0.001, TMT: t(40) = 2.55, p = 0.019) as well as to peanut oil in Veh controls (Peanut oil *vs* TMA: t(60) = 4.50, q < 0.001; Peanut oil *vs* TMT: t(60) = 2.58, q = 0.019), and this effect is accentuated in CORT mice (Group: F(1,20) = 12.78, p = 0.002; Odor: F(3,60) = 56.24, p < 0.001; Interaction: F(3,60) = 3.09, p = 0.034; ***p < 0.001). Presentation of the appetitive female urine in Veh control mice increased the distance moved compared to the peanut oil (Supplementary Figure S2D, t(60) = 2.28, q = 0.026), suggesting increased locomotion as a marker of positive olfactory valence. Bars are mean ± sem. (**E**) CORT mice approach less ♀ urine, TMA and TMT than Veh controls (Group: F(1,20) = 17.98, p < 0.001, Odor: F(3,60) = 150.80, p < 0.001, Interaction: F(3,60) = 3.03, p = 0.036; n = 11).

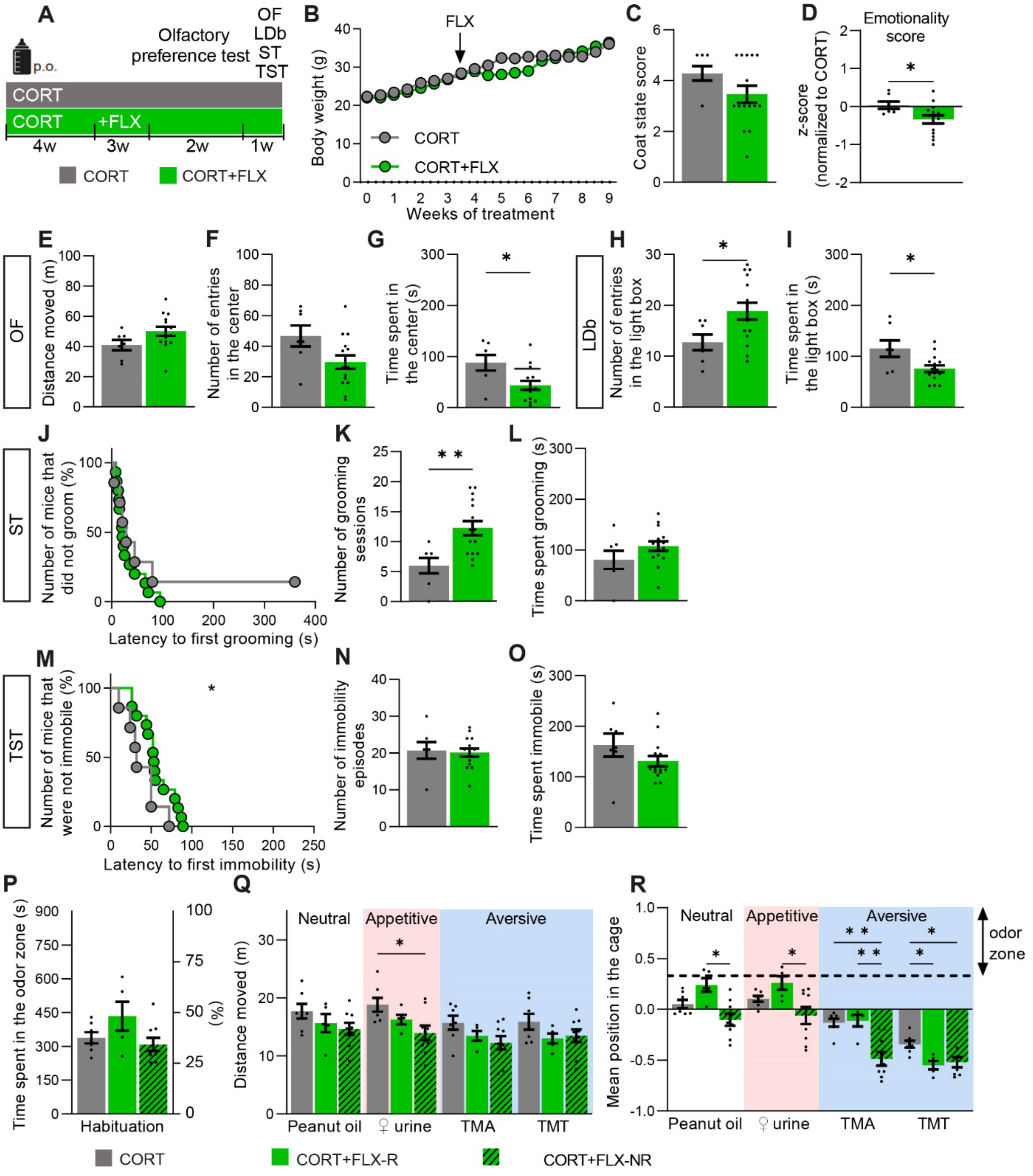


**Figure S3.** Effect of Fluoxetine on CORT phenotype. (**A**) Mice received chronic CORT to model depression (grey, n = 7), or CORT for 4 weeks then fluoxetine (FLX) in addition to CORT for the following weeks (green, n

= 14). (**B**) Body weight evolution was similar between CORT and CORT+FLX mice (Group: $F(1,20) = 1.43$ , $p = 0.246$; Weeks: $F(1.82,36.34) = 111.6$, $p < 0.0001$; Interaction: $F(18,360) = 3.94$, $p < 0.0001$). (**C**) The coat state was altered in both CORT and CORT+FLX mice ($U = 33$, $p = 0.177$). (**D**) FLX treatment decreased the global emotionality score compared to CORT ($U = 20$, $p = 0.031$). (**E-G**) CORT+Flux mice moved identically in the OF compared to CORT group (**E**: $U = 26$, $p = 0.066$; **F**: $U = 27.5$, $p = 0.080$) but explore differently the center of the arena compared to CORT (**G**: $U = 19$, $p = 0.017$). (**H-I**) FLX had opposite anxiolytic- and anxiogenic-like effects in the LDb compared to CORT (**H**: $U = 24$, $p = 0.044$; **I**: $U = 22$, $p = 0.032$). (**J-O**) Compared to CORT, FLX had an anxiolytic-like effect by increasing the number of grooming sessions in the ST (**J**: $\chi^2(1) = 0.52$, $p = 0.469$; **K**: $U = 15$, $p = 0.006$; **L**: $U = 30.5$, $p = 0.126$), and reducing the latency to first immobility in the TST (**M**: $\chi^2(1) = 4.34$, $p = 0.037$; **N**: $U = 48$, $p = 0.768$; **O**: $U = 27.5$, $p = 0.081$). (**P**) FLX treatment had no effect on the time spent in the object zone during habituation compared to CORT ($F = 3.31$, $p = 0.197$). (**Q**) CORT+FLX-NR mice moved less upon presentation of female urine than CORT mice (Group: $F(2,18) = 3.27$, $p = 0.061$; Odor: $F(2.33,42.09) = 8.58$, $p = 0.0004$; Interaction: $F(6,54) = 0.794$, $p = 0.578$). (**R**) CORT+FLX-R mice explore more peanut oil, female urine, but not TMA, than CORT+FLX-NR mice. Surprisingly, CORT+FLX-NR mice explore less TMA and both CORT+FLX-R and NR increase avoidance to TMT (Group: $F(2,18) = 9.63$, $p = 0.001$; Odor: $F(2.28,41.06) = 106.1$, $p < 0.0001$; Interaction: $F(6,54) = 5.81$, $p = 0.0001$).

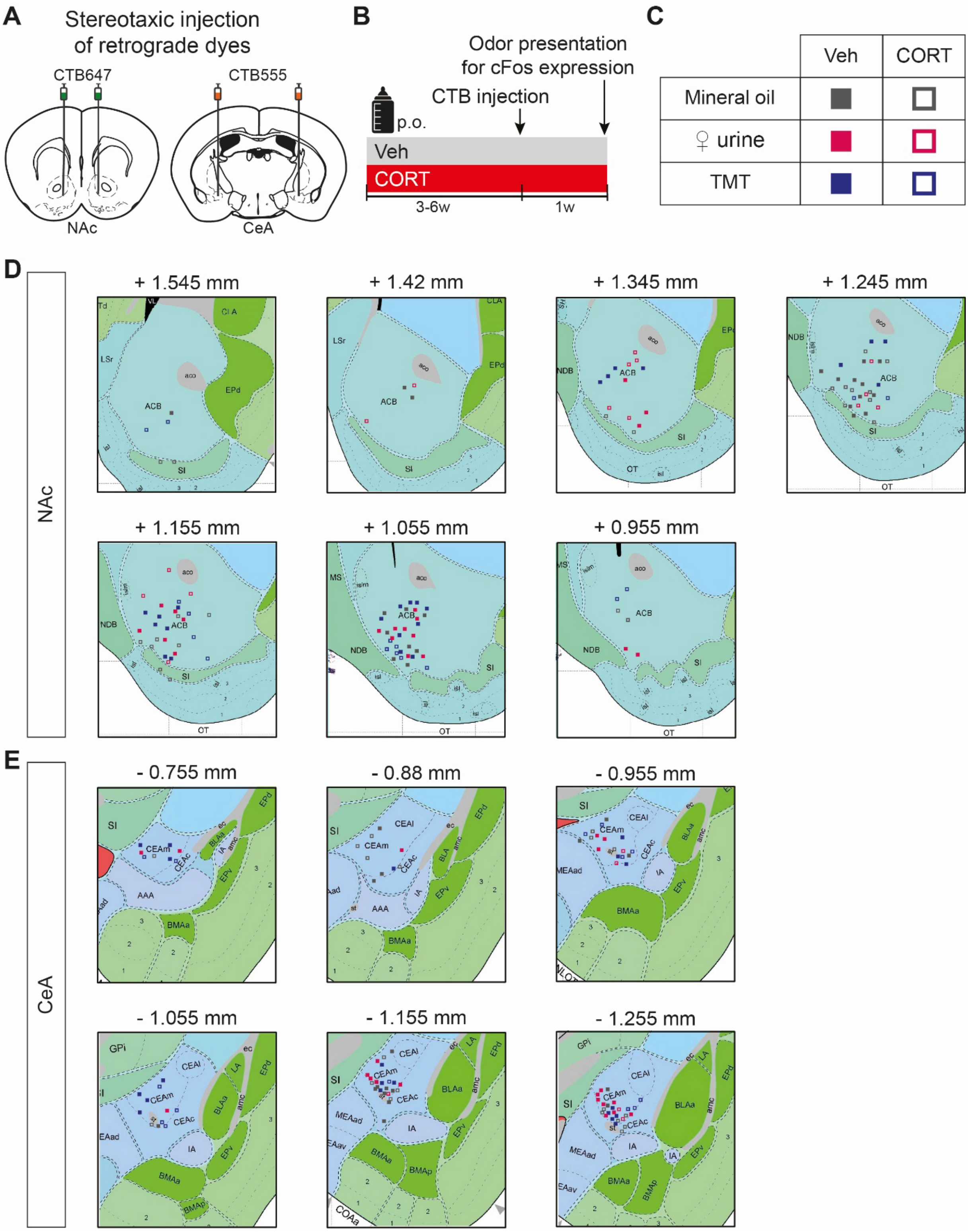


**Figure S4.** Location of stereotaxic CTB injection sites. (**A**-**B**) Retrograde fluorescent CTB dyes were injected in the NAc and the CeA before presenting odors to trigger cFos expression. (**C**) Table presenting the symbol code used in (**D**-**E**). (**D**) Injection sites for the NAc (Hemispheres, n = 116; Mice, n = 66). (**E**) Injection sites for the CeA (Hemispheres, n = 95; Mice, n = 66).

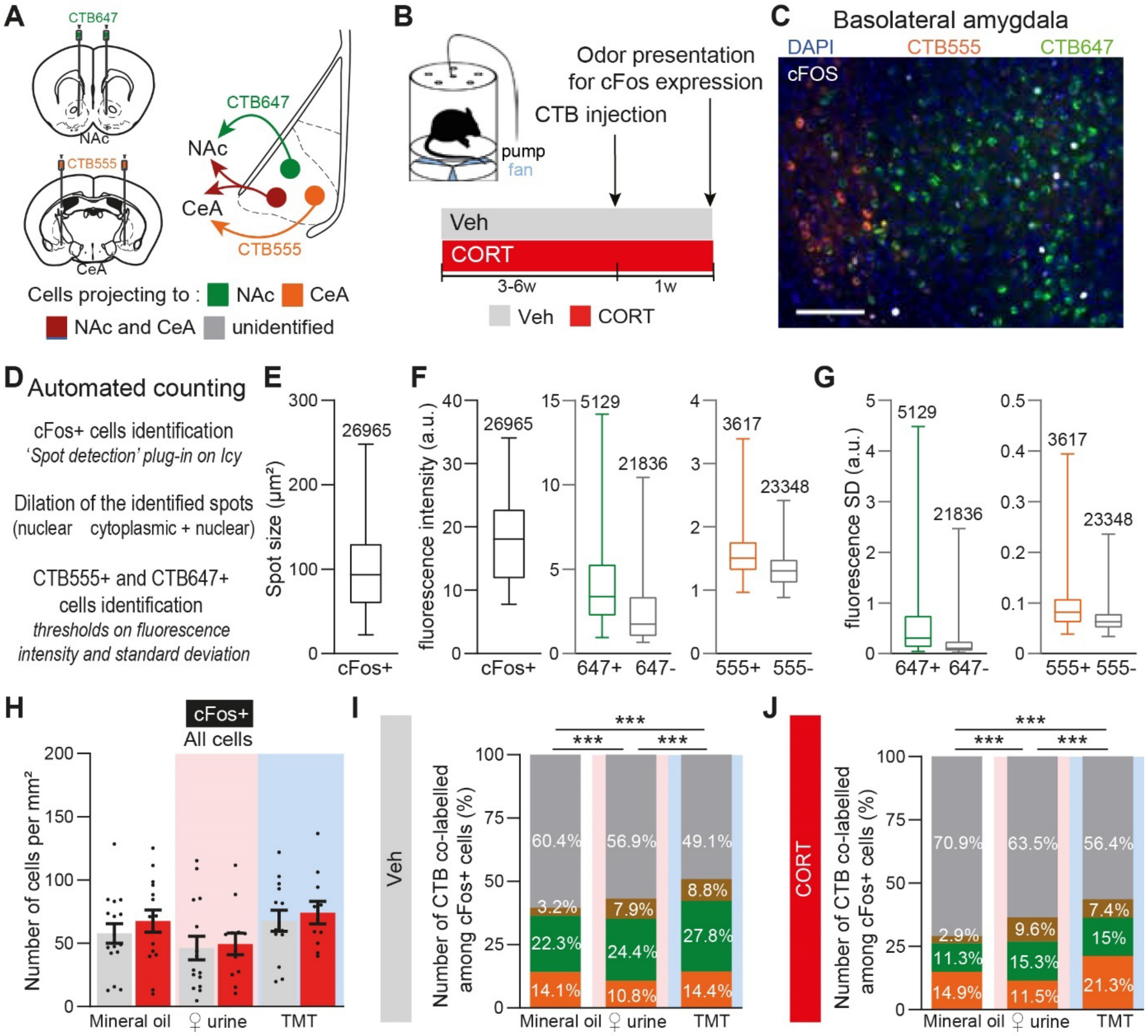


**Figure S5.** Differential BLA circuits activation in response to appetitive and aversive odors in Veh and CORT groups (**A**-**B**) Retrograde fluorescent CTB647 and CTB555 dyes were injected in the NAc (green) and the CeA (orange) respectively. Appetitive (pink), neutral (dark grey) and aversive (blue) odors were presented to trigger the immediate-early gene cFos expression in CORT and Veh mice. (**C**) Representative image of BLA cFos expression colocalized with CTB647 and/or CTB555. Scale bar, 100 µm. (**D**) Scheme of automated cell counting protocol. (**E**-**G**) Box-and-whiskers plots of the size (**e**) and fluorescence intensity (**F** left) of identified and dilated cFos+ spots, which were then sorted depending on their far red (647+/647-) or red (555+/555-) fluorescence intensity (**F** middle and right) and standard deviation (SD, **G**). (**H**) The density of cFos+ cells was similar across groups and odors (Group: $F(1,77) = 0.82$, $p = 0.368$; Odor: $F(2,77) = 3.45$, $p = 0.037$; Interaction: $F(2,77) = 0.07$, $p = 0.930$, n = 12-16; post-hoc analyses with FDR correction for the Odor effect were performed on each group separately and no statistically significant difference was found). (**I**-**J**) Proportion of activated BLA cells projecting to the NAc, the CeA or to both structures in Veh (**I**) and CORT (**J**) groups changes upon presentation of odorless mineral oil (1459 and 2777 cells corresponding to 8 and 9 mice respectively), ♀urine (1574 and 417 cells corresponding to 9 and 3 mice respectively) and TMT (3575 and 1749 cells corresponding to 12 and 7 respectively) (**I**, $\chi^2(6) = 98.44$, $p < 0.001$; Mineral oil vs ♀urine: $\chi^2(3) = 39.54$, $p < 0.001$; Mineral oil vs TMT: $\chi^2(3) = 81.77$, $p < 0.001$; ♀urine vs TMT: $\chi^2(3) = 29.11$, $p < 0.001$; **J**, $\chi^2(6) = 142.60$, $p < 0.001$; Mineral oil vs ♀urine: $\chi^2(3) = 53.29$, $p < 0.001$; Mineral oil vs TMT: $\chi^2(3) = 117.40$, $p < 0.001$; ♀urine vs TMT: $\chi^2(3) = 21.69$, $p < 0.001$). ***p < 0.001. Bars are mean ± sem.; a.u. : arbitrary unit.

| Type of cells | Veh | | | CORT | | |
|---|---|---|---|---|---|---|
| | Mineral oil | ♀urine | TMT | Mineral oil | ♀urine | TMT |
| cFos+/CTB647+ | 12.15 ± 2.35 (12) | 9.43 ± 2.45 (10) | 19.51 ± 3.37 (13) | 5.99 ± 2.31 (10) | 10.43 ± 3.11 (9) | 14.80 ± 4.40 (10) |
| Group: $F(1,58) = 1.67$, $p = 0.202$; Odor: $F(2,58) = 4.22$, p = **0.020**; Interaction: $F(2,58) = 0.70$, $p = 0.502$ | | | | | | |
| Veh ♀urine *vs* Veh TMT: $t = 2.365$, $q = 0.064$ | | | | | | |
| cFos+/CTB555+ | 10.33 ± 2.07 (13) | 6.13 ± 1.28 (13) | 10.94 ± 1.72 (13) | 13.58 ± 2.10 (13) | 13.94 ± 5.43 (5) | 16.50 ± 3.71 (9) |
| Group: $F(1,60) = 7.33$, p = **0.009**; Odor: $F(2,60) = 0.97$, $p = 0.386$; Interaction: $F(2,60) = 0.41$, $p = 0.667$ | | | | | | |
| cFos/CTB647+/ CTB555+ | 0.73 ± 0.29 (7) | 1.76 ± 0.54 (8) | 6.39 ± 1.20 (12) | 1.77 ± 0.73 (9) | 4.27 ± 2.76 (3) | 4.72 ± 2.22 (7) |
| Group: $F(1,40) = 0.30$, $p = 0.588$; Odor: $F(2,40) = 6.22$, p = **0.005**; Interaction: $F(2,40) = 1.19$, $p = 0.314$ | | | | | | |
| Veh Mineral oil *vs* Veh TMT: $t = 3.36$, q = **0.005**<br>Veh ♀urine vs Veh TMT: $t = 2.87$, q = **0.010** | | | | | | |

**Table S1.** The density of cFos+/CTB647+, cFos/CTB555+ and CTB647+/555+ cells (calculated as number of cells per mm$^2$) in the BLA is mostly similar across groups and odors. Data are reported as mean ± sem (n). Two-way ANOVA statistical analysis was followed by post-hoc FDR comparisons performed either on each group separately when the main Odor effect was statistically significant, or on each odor separately when the main Group effect was statistically significant. Only post-hoc comparisons with statistical trend or significance are indicated here.

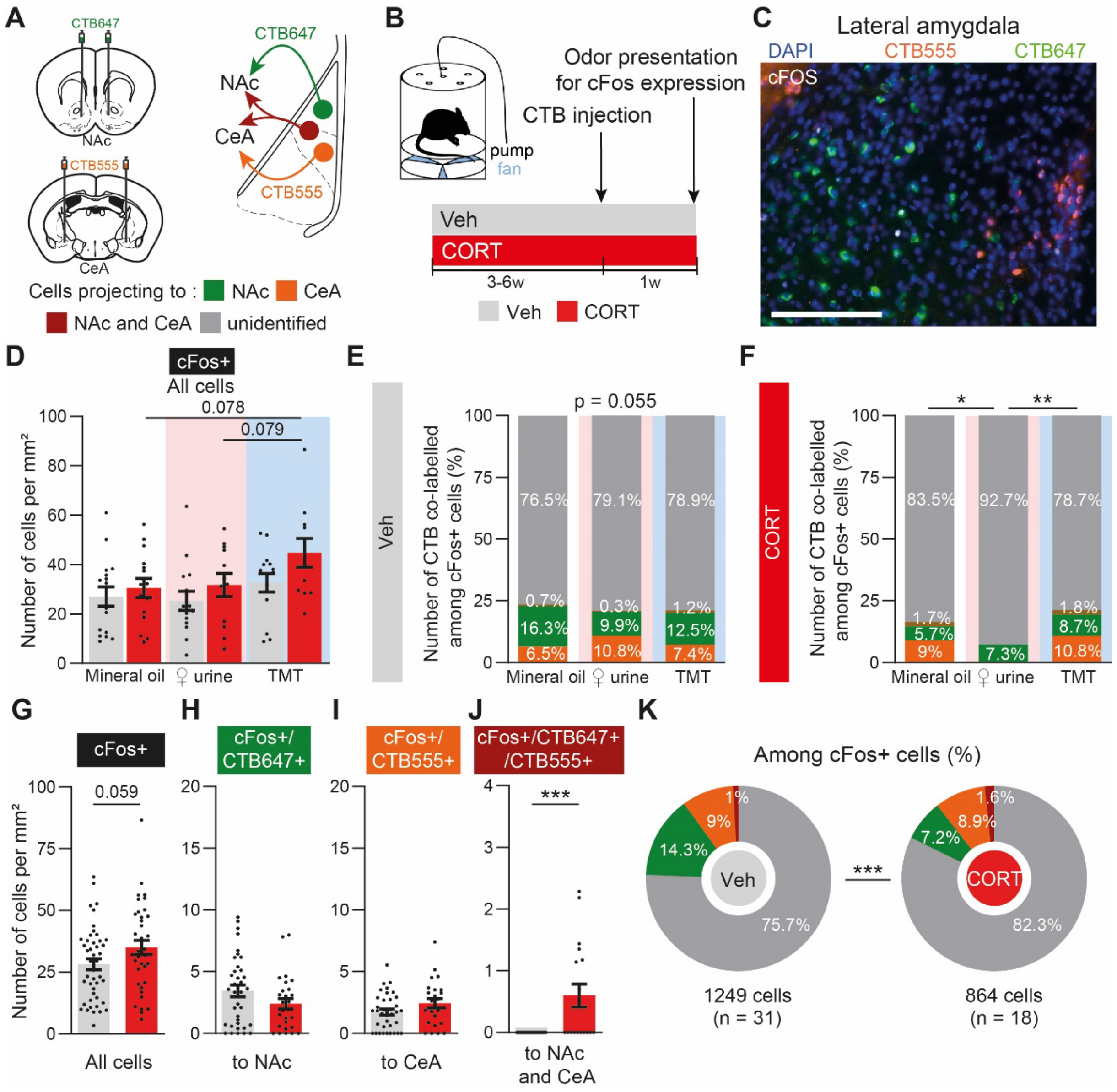


**Figure S6.** LA circuits activity upon presentation of appetitive and aversive odors in Veh and CORT mice. (**A-B**) Retrograde fluorescent CTB647 and CTB555 dyes were injected in the NAc (green) and the CeA (orange) respectively. Appetitive (pink), neutral (dark grey) and aversive (blue) odors were presented to trigger the immediate-early gene cFos expression in CORT and Veh mice. (**C**) Representative image of LA cFos expression colocalized with CTB647 and/or CTB555. Scale bar, 100 µm. (**D**) The density of cFos+ cells tended to increase in CORT mice in response to TMT compared to mineral oil and ♀urine (Group: $F(1,77) = 4.40$, $p = 0.036$; Odor: $F(2,77) = 3.47$, $p = 0.036$; Interaction: $F(2,77) = 0.52$, $p = 0.594$, n = 11-16; post-hoc analyses with FDR correction for the Odor effect were performed on each group separately and for the Group effect on each odor separately, for which no difference was found statistically significant). (**E-F**) Proportion of activated LA projecting cells to the NAc, the CeA or to both structures in Veh mice (**E**) tends to change and CORT mice (**F**) changes upon presentation of odorless mineral oil (307 and 401 cells corresponding to 8 and 8 mice respectively), ♀urine (344 and 82 cells corresponding to 9 and 3 mice respectively) and TMT (758 and 381 cells corresponding to 12 and 7 mice respectively) (**E**, $\chi^2(6) = 12.32$, $p = 0.055$; **F**, $\chi^2(6) = 14.32$, $p = 0.026$; Mineral oil vs ♀urine: $\chi^2(3) = 9.75$, $p = 0.021$; Mineral oil vs TMT: $\chi^2(3) = 3.53$, $p = 0.317$; ♀urine vs TMT: $\chi^2(3) = 12.09$, $p = 0.007$). (**G-J**) Regardless of which odor was used for cFos stimulation, CORT mice display a trend to increased density of LA cFos+ (**G**, $t(81) = 1.92$, $p = 0.059$, n = 38-45) and increased cFos+/CTB647+/CTB555+ (**J**, $U = 105$, $p < 0.001$, n = 18-21) cell density relative to Veh controls, but no difference regarding LA cFos+/CTB647+ (**H**, $U = 377.5$, $p = 0.133$, n = 27-36) and cFos+/CTB555+ (**I**, $t(58) = 1.645$, $p = 0.106$, n = 24-36) density. (**K**) Distribution of CTB647 and/or CTB555 colocalization among the total number of cFos+ cells in the LA differs between CORT and Veh groups ($\chi^2(3) = 27.61$, $p < 0.001$). *$p < 0.05$, **$p < 0.01$, ***$p < 0.001$. Bars are mean ± sem.

<table>
<tr><td rowspan="2">Type of cells</td><td colspan="3">Veh</td><td colspan="3">CORT</td></tr>
<tr><td>Mineral oil</td><td>♀urine</td><td>TMT</td><td>Mineral oil</td><td>♀urine</td><td>TMT</td></tr>
<tr><td>cFos+/CTB647+</td><td>2.78 ± 1.03<br>(12)</td><td>2.91 ± 0.74<br>(11)</td><td>4.55 ± 0.62<br>(13)</td><td>1.25 ± 0.52<br>(9)</td><td>2.90 ± 0.74<br>(9)</td><td>4.52 ± 1.64<br>(10)</td></tr>
<tr><td colspan="7">Group: F(1,58) = 0.45, p = 0.507; Odor: F(2,58) = 3.68, p = 0.031; Interaction: F(2,58) = 0.41, p = 0.667</td></tr>
<tr><td colspan="7">CORT Mineral oil vs CORT TMT: t = 2.29, q = 0.077</td></tr>
<tr><td>cFos+/CTB555+</td><td>1.43 ± 0.34<br>(12)</td><td>1.98 ± 0.43<br>(13)</td><td>1.80 ± 0.52<br>(11)</td><td>2.60 ± 0.33<br>(12)</td><td>1.44 ± 1.01<br>(5)</td><td>4.96 ± 1.55<br>(9)</td></tr>
<tr><td colspan="7">Group: F(1,58) = 4.47, p = 0.039; Odor: F(2,56) = 2,95, p = 0.061; Interaction: F(2,56) = 2.91, p = 0.063</td></tr>
<tr><td colspan="7">Veh TMT vs CORT TMT: t = 3.14, p = 0.008</td></tr>
<tr><td>cFos/CTB647+/<br>CTB555+</td><td>0.00 ± 0.00<br>(6)</td><td>0.00 ± 0.00<br>(8)</td><td>0.31 ± 0.15<br>(11)</td><td>0.67 ± 0.30<br>(8)</td><td>0.00 ± 0.00<br>(3)</td><td>0.78 ± 0.32<br>(7)</td></tr>
<tr><td colspan="7">Group: F(1,37) = 4.00, p = 0.053; Odor: F(2,37) = 2.65, p = 0.083; Interaction: F(2,37) = 0.92, p = 0.409</td></tr>
</table>

**Table S2.** The density of cFos+/CTB647+, cFos/CTB555+ and CTB647+/555+ cells (calculated as a number of cells per $mm^2$) in the LA is mostly similar across groups and odors. Data are reported as mean ± sem (n). Two-way ANOVA statistical analysis was followed by post-hoc FDR comparisons performed either on each group separately when the main Odor effect was statistically significant, or on each odor separately when the main Group effect was statistically significant. Only post-hoc comparisons with statistical trend or significance are indicated here.

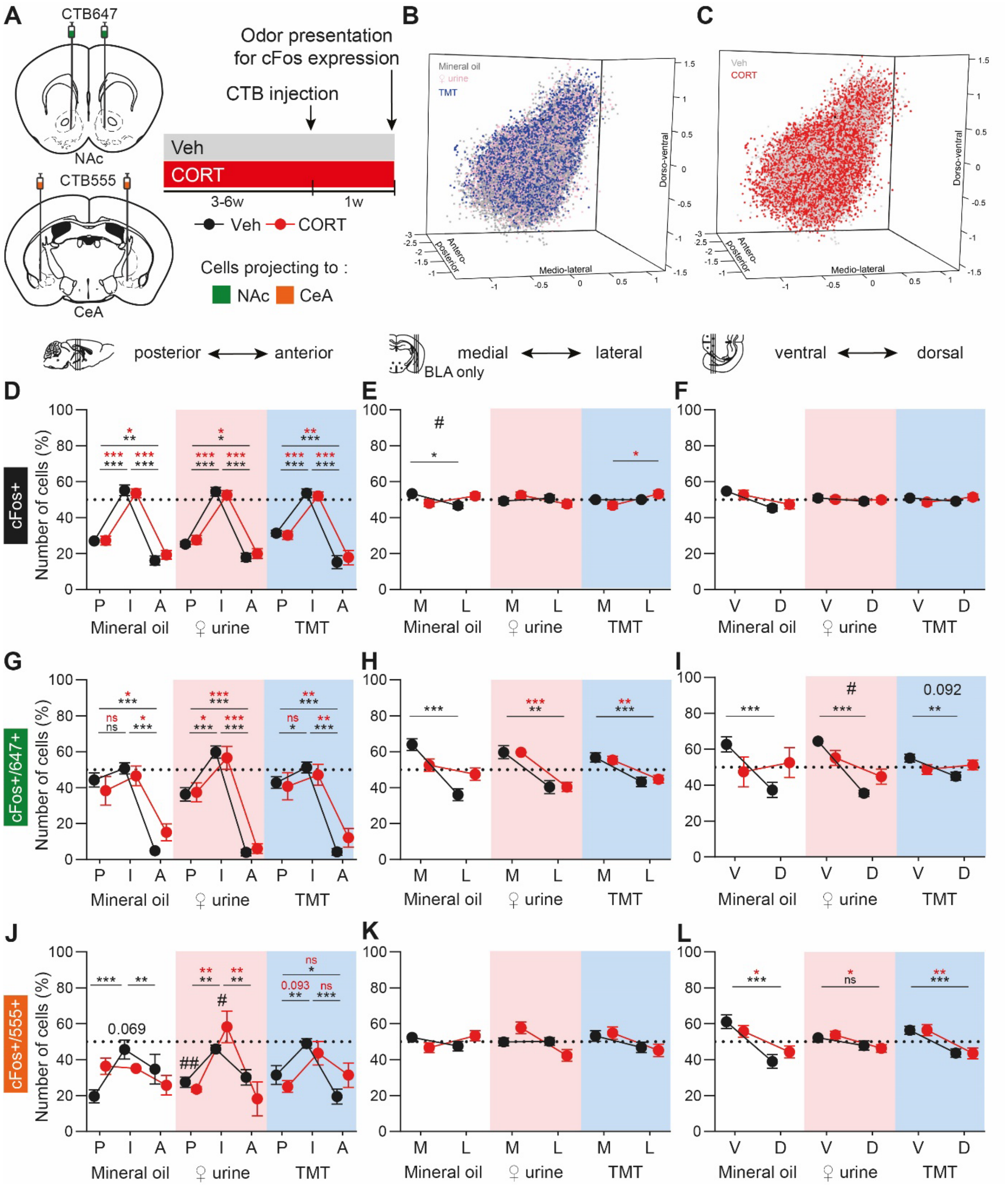


**Figure S7.** Spatial location of cFos+ and colocalizing cFos+/CTB+ cells through LA/BLA in response to appetitive and aversive odors in both Veh and CORT mice. (**A**) Odorless mineral oil, appetitive ♀urine and aversive TMT were presented to stimulate the immediate-early gene cFos expression, one week after stereotaxic injection of retrograde dyes CTB647 and CTB555 in the NAc and the CeA (centromedial part) respectively, in chronically CORT or Veh-treated mice. (**B**-**C**) 3-dimensional plot of all the cFos+ cells responding to different odors (**B**) in the different groups (**C**) (n = 11-16, corresponding to 32,288 cFos+ cells). (**D**-**L**) Spatial distribution of cFos+ (**D**-**F**), cFos+/CTB647+ (**G**-**I**) and cFos+/CTB555+ (**J**-**L**) cells along the antero-posterior (**D**,**G**,**J**), medio-lateral (**E**,**H**,**K**) and dorso-ventral (**F**,**I**,**L**) axis. Along the antero-posterior axis, the anterior part (A) corresponds to [-0.6;-1.4[ mm respective to bregma, the intermediate part (I) to [-1.4;-2[ mm and the posterior part (P) to [-2;-3] mm. For the medio-lateral axis, only BLA cells were included. The medial part (M) corresponds to [-1.4;-0.2] mm respective to the center of the drawn region of interest (ROI) around the LA/BLA while the lateral part (L) corresponds to ]-0.2;1.4] mm to the ROI center. Concerning the dorso-ventral axis, the cut was made around the BLA/LA separation. The ventral part (V) corresponds to [-1.6;-0.2] mm to the ROI center, while the dorsal part corresponds to ]-0.2;1.6] mm to the ROI center. Three-way repeated measures ANOVA analyses are reported in **Table S3**. Black and red stars (*) represent post-hoc statistically significant differences along axes for the Veh and CORT groups respectively. Hashtags (#) represent post-hoc statistically significant differences between Veh and CORT, also reported in **Table S3**. $^{*/\#}p < 0.05$, $^{**/\#\#}p < 0.01$, $^{***}p < 0.001$, ns: not significant. Data are points ± sem.

| Type of cell / Axis | cFos+ | | cFos+/CTB647+ | | cFos+/CTB555+ | |
|---|---|---|---|---|---|---|
| **Antero-posterior** | | p-value | | p-value | | p-value |
| Axis | F (1.67,125.52) = 221.16 | **<0.001** | F (1.43,77.19) = 95.85 | **<0.001** | F (1.68,92.6) = 23.64 | **<0.001** |
| Odor | F (2,75) = 0.00 | 1.000 | F (2,54) = 0.00 | 1.000 | F (2,55) = 0.00 | 1.000 |
| Group | F (1,75) = 0.00 | 1.000 | F (1,54) = 0.00 | 1.000 | F (1,55) = 0.00 | 1.000 |
| Axis x Odor | F (3.35,125.52) = 0.76 | 0.531 | F (2.86,77.19) = 1.78 | 0.161 | F (3.37,92.6) = 0.94 | 0.434 |
| Axis x Group | F (1.67,125.52) = 1.08 | 0.332 | F (1.43,77.19) = 1.64 | 0.206 | F (1.68,92.6) = 0.22 | 0.762 |
| Odor x Group | F (2,75) = 0.00 | 1.000 | F (2,54) = 0.00 | 1.000 | F (2,55) = 0.00 | 1.000 |
| Axis x Odor x Group | F (3.35,125.52) = 0.07 | 0.98 | F (2.86,77.19) = 0.20 | 0.890 | F (3.37,92.6) = 3.08 | **0.026** |
| cFos+/CTB555+; Veh MO I vs CORT MO I: t = 3.19, q = **0.005**<br>Veh MO P vs CORT MO P: t = 2.60, q = **0.017**<br>Veh FU I vs CORT FU I: t = 1.96, q = *0.069*<br>Veh FU A vs Veh TMT A: t = 1.78, q = *0.087*<br>CORT MO I vs CORT FU I: t = 3.26, q = **0.006**<br>CORT MO P vs CORT FU P: t = 1.80, q = *0.092*<br>CORT MO P vs CORT TMT P: t = 1.92, q = *0.070* | | | | | | |
| **Medio-lateral** | p-value | | p-value | | | p-value |
| Axis | F (1,77) = 0.00 | 0.952 | F (1,53) = 39.72 | **<0.001** | F (1,60) = 3.89 | *0.053* |
| Odor | F (2,77) = 0.00 | 1.000 | F (2,53) = 0.00 | 1.000 | F (2,60) = 0.00 | 1.000 |
| Group | F (1,77) = 0.00 | 1.000 | F (1,53) = 0.00 | 1.000 | F (1,60) = 0.00 | 1.000 |
| Axis x Odor | F (2,77) = 1.08 | 0.344 | F (2,53) = 0.71 | 0.498 | F (2,60) = 1.53 | 0.225 |
| Axis x Group | F (1,77) = 1.52 | 0.222 | F (1,53) = 2.90 | *0.095* | F (1,60) = 0.25 | 0.622 |
| Odor x Group | F (2,77) = 0.00 | 1.000 | F (2,53) = 0.00 | 1.000 | F (2,60) = 0.00 | 1.000 |
| Axis x Odor x Group | F (2,77) = 3.17 | **0.048** | F (2,53) = 1.95 | 0.152 | F (2,60) = 2.47 | *0.094* |
| cFos+; Veh MO M *vs* CORT MO M: t = 2.07, q = **0.047** | | | | | | |
| **Dorso-ventral** | p-value | | p-value | | p-value | |
| Axis | F (1,77) = 3.15 | *0.080* | F (1,54) = 9.19 | **0.004** | F (1,60) = 18.66 | **<0.001** |
| Odor | F (2,77) = 0.00 | 1.000 | F (2,54) = 0.00 | 1.000 | F (2,60) = 0.00 | 1.000 |
| Group | F (1,77) = 0.00 | 1.000 | F (1,54) = 0.00 | 1.000 | F (1,60) = 0.00 | 1.000 |
| Axis x Odor | F (2,77) = 2.66 | *0.077* | F (2,54) = 1.54 | 0.225 | F (2,60) = 1.24 | 0.296 |
| Axis x Group | F (1,77) = 1.25 | 0.267 | F (1,54) = 7.78 | **0.007** | F (1,60) = 0.20 | 0.656 |
| Odor x Group | F (2,77) = 0.00 | 1.000 | F (2,54) = 0.00 | 1.000 | F (2,60) = 0.00 | 1.000 |
| Axis x Odor x Group | F (2,77) = 0.11 | 0.899 | F (2,54) = 0.52 | 0.596 | F (2,60) = 0.65 | 0.525 |
| cFos+/CTB647+; Veh FU V vs CORT FU V: t = 2.13, q = **0.048**<br>Veh TMT V vs CORT TMT V: t = 1.78, q = *0.092* | | | | | | |

**Table S3.** Corresponding statistical analyses for **Figure S6** (Three-way repeated measures ANOVA or Mixed-effect model). For the medio-lateral and dorso-ventral axes, post-hoc analyses comparing Veh and CORT are reported for the medial part (M) and ventral part (V), but they have respectively identical adjusted p-values (q) than the lateral and dorsal parts. A: anterior part, I: intermediate part, P: posterior part.

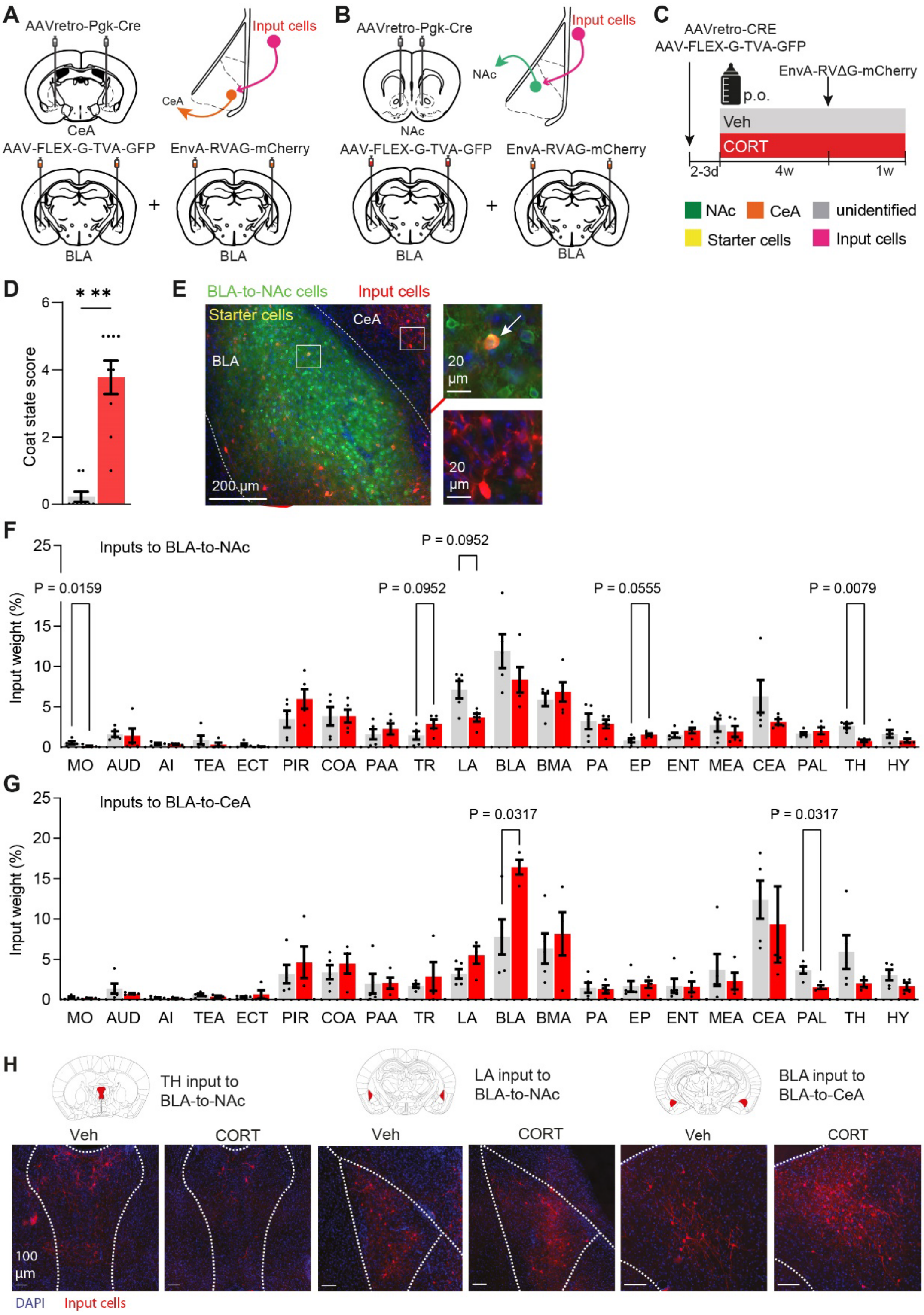


**Figure S8.** CORT treatment changes synaptic connectivity of BLA neurons. (**A**) AAVr-Pgk-Cre in the NAc and AAV-hSyn-Flex-G-TVA-GFP and EnvA-RVAG-mCherryin the BLA were injected to trace monosynaptic input to BLA-to-CeA projecting cells (in orange, n = 9). (**B**) Same than A except that AAVr-Pgk-Cre were injected in the CeA to labelled BLA-to-NAc cells (in green, n = 9). (**C**) Mice received vehicle (grey, n = 10) or chronic CORT (red, n = 9) to model depression. (**D**)The coat state was significantly altered in CORT mice (U = 1, p = 0.0001). (**E**) Representative image of a Veh mouse in which we labelled BLA-to-NAc cells (green). Input cells are labelled in red, and starter cells (yellow) are defined by the colocalization of green and red markers. An example starter cell is pointed by an arrow in the top right panel. (**F-G**) Extended quantification of BLA-to-NAc (**F**) and BLA-to-CeA (**G**) cells inputs in Veh and CORT mice [Mann-Whitney tests, n=4-5; total cells analysed 26639 (F) and 24255 (G)]. (**H**) Representative images of TH inputs to BLA-to-NAc cells, LA inputs to BLA-to-

NAc cells, and BLA inputs to BLA-to-CeA cells in a Veh and CORT mice. Input cells: red; DAPI: blue. Scale bars, 20 µm. Bars are mean ± sem. ***p < 0.001.

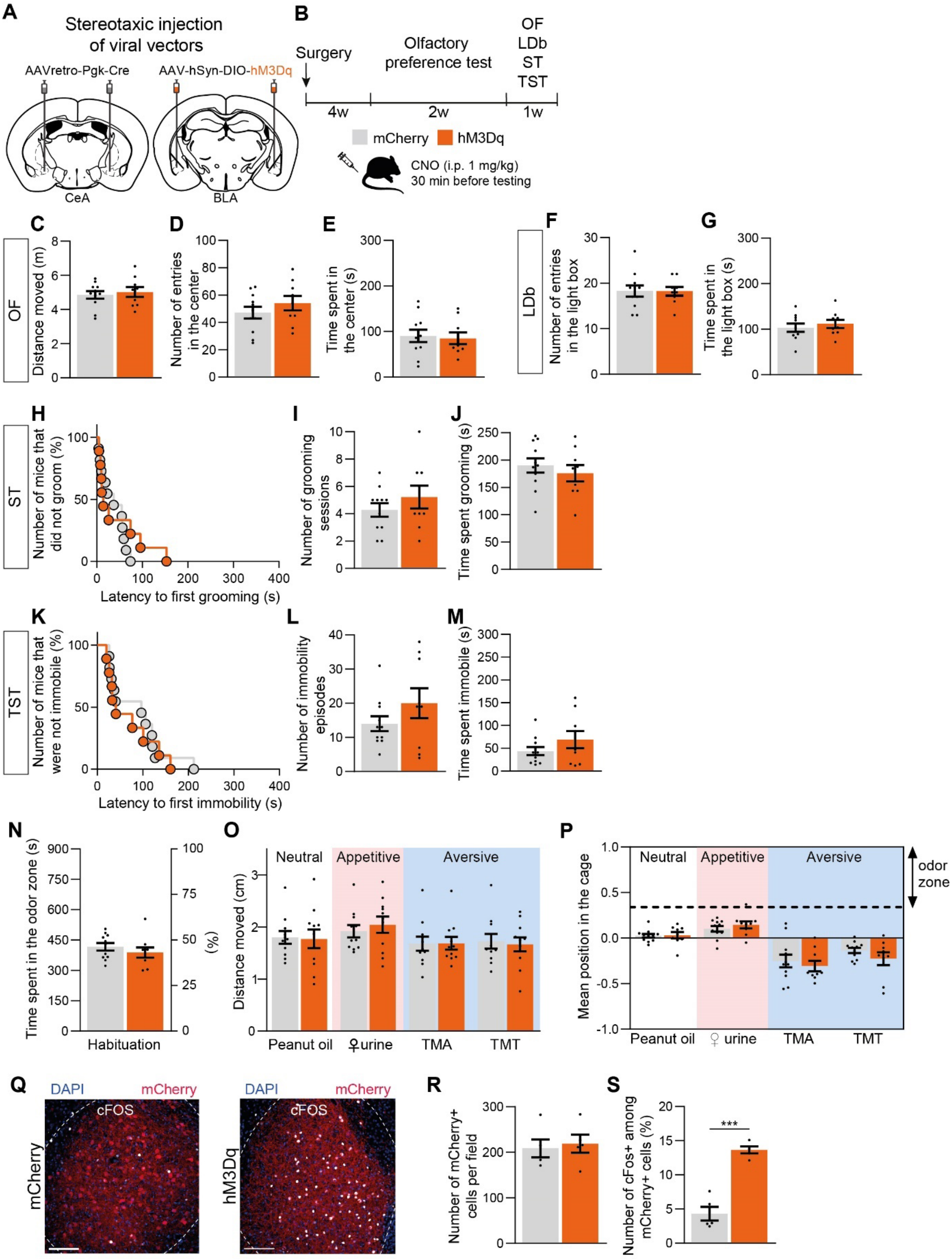


**Figure S9.** Chemogenetic BLA-to-CeA cells activation does not promote anxiety- and depressive-like behaviors. (**A**-**B**) AAVr-Pgk-Cre in the CeA and AAV-hSyn-DIO-hM3Dq-mCherry (orange, n = 9, or AAV-hSyn-DIO-mCherry for the controls, grey, n = 11) in the BLA were injected to activate BLA-to-CeA cells. (**C**-**E**) CNO injection had no effect in the OF in hM3Dq mice relative to mCherry controls (**C**, $t(18) = 0.46$, $p = 0.650$; **D**, $t(18) = 1.02$, $p = 0.321$; **E**, $t(18) = 0.28$, $p = 0.785$). (**F**-**G**) hM3Dq mice were not different than mCherry controls in the LDb (**F**, $t(18) = 0.03$, $p = 0.098$; **G**, $t(17) = 0.65$, $p = 0.522$, n = 10-11) and ST (**H**, $\chi^2(1) = 0.46$, $p = 0.500$; **I**, $t(18) = 1.03$, $p = 0.317$; **J**, $t(18) = 0.73$, $p = 0.476$). (**K**-**M**) CNO injection did not change immobility behavior of hM3Dq mice compared to mCherry controls in the TST (**K**, $\chi^2(1) = 0.19$, $p = 0.667$; **L**, $t(18) = 1.30$, $p = 0.209$; **M**, $t(18) = 1.28$, $p = 0.216$). The global score do not difere between groups neither in the OF (**C-D**, $t(18) = 0.54$, $p = 0.598$), LDb (**F-G**, $t(18) = 0.35$, $p = 0.733$), ST (**H-J**, $t(18) = 0.02$, $p = 0.987$) nor TST (**K-M**, $t(18) = 1.24$, $p = 0.232$). **(N**) CNO injection in hM3Dq mice did not modify the time spent in the "object" zone during habituation without odor compared to mCherry controls ($t(18) = 0.93$, $p = 0.366$). (**O**) hM3Dq mice moved similarly than mCherry controls (Group: $F(1,18) = 0.05$, $p = 0.827$, Odor: $F(3,54) = 9.34$, $p < 0.001$; Interaction: $F(3,54) = 0.91$, $p = 0.444$). (**P**) Chemogenetic BLA-to-CeA cells activation did not affect the behavior in the olfactory preference test (Group: $F(1,18) = 0.28$, $p = 0.605$; Odor: $F(3,54) = 43.65$, $p < 0.001$; Interaction: $F(3,54) = 1.15$, $p = 0.338$)). (**Q**) Representative images of BLA cFos expression. Scale bar, 125 µm. (**R**-**S**) Quantification of the mCherry+ cell number in mCherry and hM3Dq mice (**R**, $t(8) = 0.37$, $p = 0.722$) and the percentage of cFos expression among mCherry+ cells after CNO injection (**S**, $t(8) = 8.31$, ***$p < 0.001$). Bars are mean ± sem.

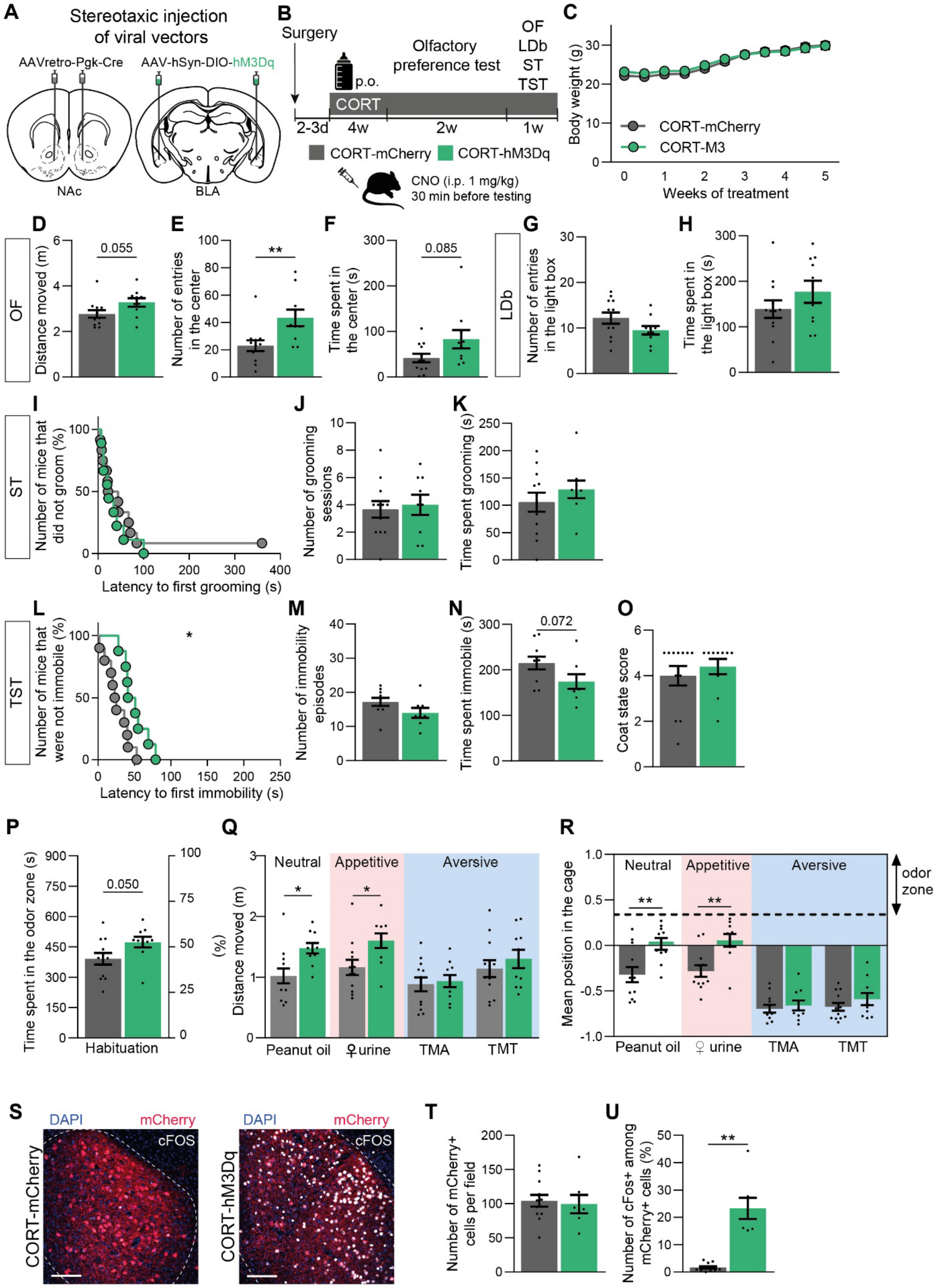
A
Stereotaxic injection of viral vectors
AAVretro-Pgk-Cre
AAV-hSyn-DIO-hM3Dq
NAc
BLA
B
Surgery
p.o.
Olfactory preference test
OF
LDb
ST
TST
CORT
2-3d
4w
2w
1w
CORT-mCherry
CORT-hM3Dq
CNO (i.p. 1 mg/kg)
30 min before testing
C
Body weight (g)
CORT-mCherry
CORT-M3
Weeks of treatment
D
OF
Distance moved (m)
0.055
E
Number of entries in the center
**
F
Time spent in the center (s)
0.085
G
LDb
Number of entries in the light box
H
Time spent in the light box (s)
I
ST
Number of mice that did not groom (%)
Latency to first grooming (s)
J
Number of grooming sessions
K
Time spent grooming (s)
L
TST
Number of mice that were not immobile (%)
*
Latency to first immobility (s)
M
Number of immobility episodes
N
Time spent immobile (s)
0.072
O
Coat state score
P
Time spent in the odor zone (s)
(%)
0.050
Habituation
Q
Neutral
Appetitive
Aversive
Distance moved (m)
*
Peanut oil
♀urine
TMA
TMT
R
Neutral
Appetitive
Aversive
odor zone
Mean position in the cage
**
Peanut oil
♀ urine
TMA
TMT
S
DAPI
mCherry
cFOS
CORT-mCherry
CORT-hM3Dq
T
Number of mCherry+ cells per field
U
Number of cFos+ among mCherry+ cells (%)
**

**Figure S10.** Chemogenetic BLA-to-NAc cells activation has anxiolytic and antidepressant-like effects on CORT mice. (**A**-**B**) AAVr-Pgk-Cre in the NAc and AAV-hSyn-DIO-hM3Dq-mCherry (green, or AAV-hSyn-DIO-mCherry for the controls. grey) in the BLA were injected to activate BLA-to-NAc cells. (**C**) Body weight evolution was similar between CORT-mCherry and CORT-hM3Dq mice (Group: $F(1,20) = 0.12$, $p = 0.736$; Weeks: $F(10,200) = 181.80$, $p < 0.001$; Interaction: $F(10,200) = 1.56$, $p = 0.121$). (**D**-**F**) CNO injection had anxiolytic-like effects in the OF in CORT-hM3Dq mice relative to CORT-mCherry controls (n = 10-12) (**D**, $t(20) = 2.90$, $^{**}p = 0.009$; **E**, $t(20) = 2.04$, $p = 0.055$; **F**, $t(12.81) = 1.87$, $p = 0.085$). (**G, H**) CORT-hM3Dq mice were not different than CORT-mCherry controls in the LDb (**G**, $t(20) = 1.67$, $p = 0.111$; **H**, $t(20) = 1.24$, $p = 0.228$, n = 10-12) and ST (**I**, $\chi^2(1) = 0.40$, $p = 0.529$; **J**, $t(19) = 0.35$, $p = 0.730$; **K**, $t(19) = 0.95$, $p = 0.354$, n = 9-12). (**L**-**N**) CNO injection increased the first immobility latency in CORT-hM3Dq mice compared to CORT-mCherry controls in the TST (n = 8-10) (**L**, $\chi^2(1) = 6.16$, $^{*}p = 0.013$) and tended to decrease the immobility time (**N**, $t(16) = 1.92$, $p = 0.072$). but had no effect on the immobility episodes number (**M**, $t(16) = 1.72$, $p = 0.106$). Overall, CNO i.p. injection decreased anxiety-like phenotype in CORT-hM3Dq mice in the the OF ( $t(11.46) = 3.30$, $^{**}p = 0.007$, n = 10-12) and was antidepressant on the TST ( $t(16) = 3.85$, $^{**}p = 0.001$, n = 8-10) compared to CORT-mCherry controls, but had no effect in the LDb ( $t(20) = 0.09$, $p = 0.926$, n = 10-12) or ST ( $t(19) = 0.82$, $p = 0.420$, n = 9-12) when the emotionality score for these test was analysed. (**O**) The coat state was altered in both CORT-hM3Dq and CORT-mCherry mice (U = 51.50, $p = 0.529$, n = 10-12). (**P**) CNO injection in CORT-hM3Dq mice tends to increase the time spent in the “object” zone during habituation without odor compared to CORT-mCherry controls ($t(20) = 2.09$, $p = 0.050$).(**Q**) CORT-hM3Dq mice moved more upon presentation of peanut oil and ♀urine than CORT-mCherry controls (Group: $F(1,20) = 3.20$, $p = 0.089$, Odor: $F(3,60) = 18.26$, $p < 0.001$; Interaction: $F(3,60) = 4.66$, $p = 0.005$; $^{*}p < 0.05$). (**R**) Chemogenetic BLA-to-NAc cells activation increases interest for neutral and appetitive odors in the olfactory preference test but not for aversive odors (Group: $F(1,20) = 7.37$, $p = 0.013$; Odor: $F(3,60) = 111.8$, $p < 0.001$; Interaction: $F(3,60) = 7.88$, $p < 0.001$; n = 10-12). (**S**) Representative images and magnifications of BLA cFos expression. Scale bar, 125 µm. (**T**-**U**) Quantification of the mCherry+ cell number in CORT-mCherry and CORT-hM3Dq mice (**T**, U = 36, $p = 0.650$, n = 7-12) and the percentage of cFos expression among mCherry+ cells after CNO injection (**U**, $t(6.135) = 5.51$, $^{**}p = 0.001$, n = 7-12). Bars are mean ± sem.

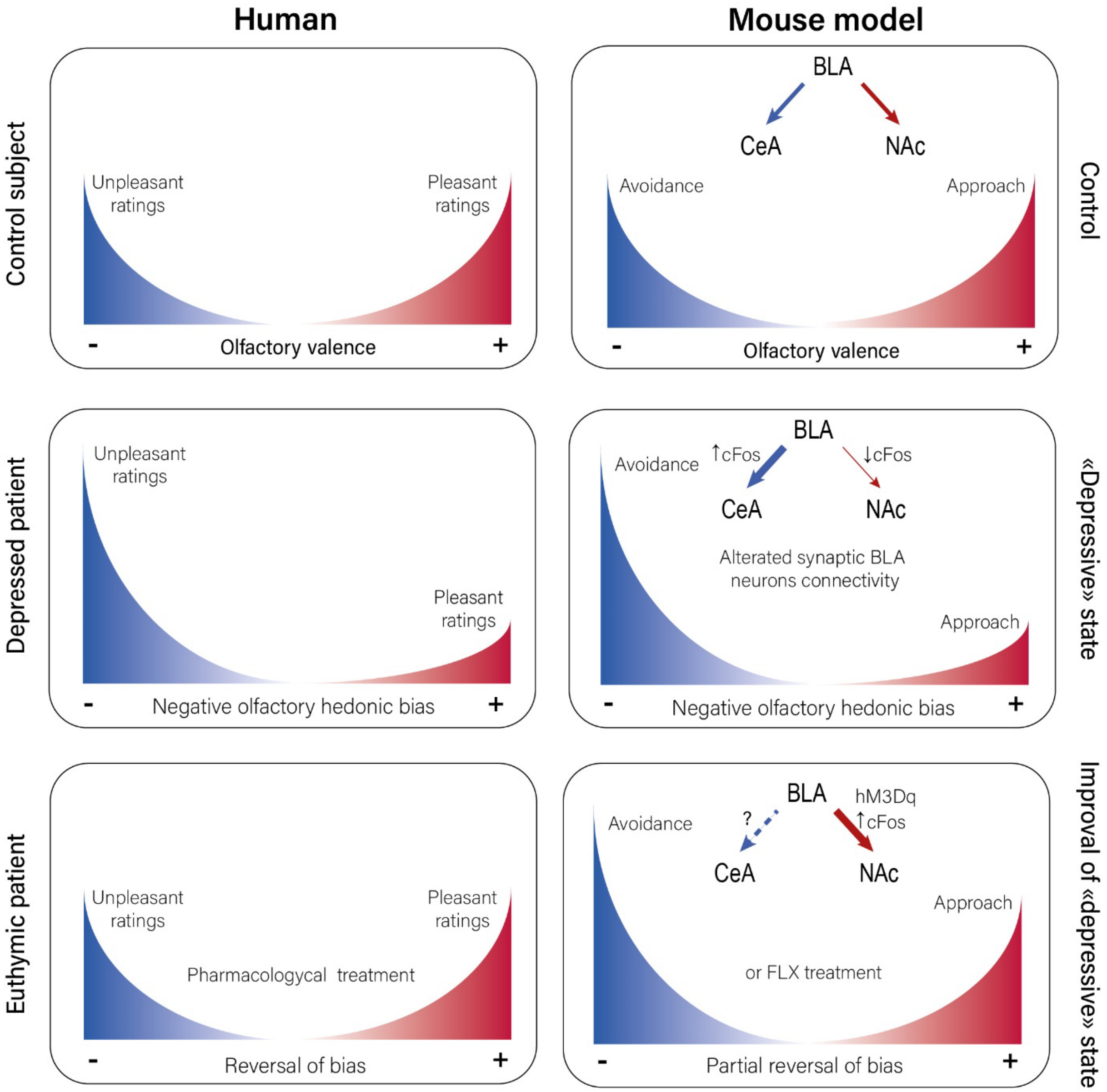


**Figure S11.** Summary scheme of the results. Depressed BD patients as well as the CORT-induced model for depression show a negative olfactory hedonic bias. Patients present decreased number of odors rated as pleasant associated with increased number of rated unpleasant odors respect to control subject, while CORT-treated mice exhibit decreased approach behavior towards appetitive odors and increased avoidance behaviour towards aversive odors respect to control group. The chronic CORT administration elevates the activity of BLA projecting neurons to the CeA, which preferentially encode negative valence, and reduces the activity of BLA projecting neurons to the NAc, preferentially encoding positive valence, as measured by the immediate early-gene cFos expression. Euthymic patients after pharmacological treatment reverse the emotional bias. In agree with these data, chemogenetic activation of the BLA-to-NAc neurons improves partially the negative hedonic olfactory bias and depressive-like phenotypes induced by chronic CORT administration, as observed in mice after responsive to fluoxetine treatment.
